# Evolution toward maximum transport capacity of the Ttg2 ABC system in *Pseudomonas aeruginosa*

**DOI:** 10.1101/834812

**Authors:** Daniel Yero, Lionel Costenaro, Oscar Conchillo-Solé, Mireia Díaz-Lobo, Adrià Mayo, Mario Ferrer-Navarro, Marta Vilaseca, Isidre Gibert, Xavier Daura

## Abstract

In *Pseudomonas aeruginosa*, Ttg2D is the soluble periplasmic phospholipid-binding component of an ABC transport system thought to be involved in maintaining the asymmetry of the outer membrane. The crystallographic structure of Ttg2D at 2.5Å resolution reveals that this protein can bind two diacyl phospholipids. Native and denaturing mass spectrometry experiments confirm that Ttg2D binds two phospholipid molecules, which may have different head groups. Analysis of the available structures of Ttg2D orthologs allowed us to classify this protein family as a novel substrate-binding protein fold and to venture the evolutionary events that differentiated the orthologs binding one or two phospholipids. In addition, gene knockout experiments in *P. aeruginosa* PAO1 and multidrug-resistant strains show that disruption of this system leads to outer membrane permeabilization. This demonstrates the role of this system in low-level intrinsic resistance against certain antibiotics that use a lipid-mediated pathway to permeate through membranes.

## Introduction

*Pseudomonas aeruginosa* are amongst the most important multidrug-resistant (MDR) human pathogens^1^, showing inherent resistance to an important number of the presently available antibiotics^2^. *P. aeruginosa* are responsible for chronic lung infections in individuals with chronic obstructive pulmonary disease or cystic fibrosis (CF)^3^ and account for over a tenth of all nosocomial infections^4^. A number of effective drugs and formulations can treat *P. aeruginosa* infections, even in CF patients^5^. These include frontline antibiotics such as piperazillin-tazobactam, ceftazidime, aztreonam, imipenem, meropenem, ciprofloxacin, levofloxacin, tobramycin, amikacin, and colistin^6^. Yet, resistance to most of these antimicrobials is being increasingly reported^7^. The basis for the inherently high resistance of these microorganisms is primarily their low outer-membrane (OM) permeability^8, 9^, complemented by the production of antibiotic-inactivating enzymes (e.g. β-lactamases), the constitutive expression of efflux pumps^10, 11^ and the capacity to form biofilms^1, 12^, among other mechanisms. The susceptibility of *P. aeruginosa* to antimicrobials can be additionally reduced by the acquisition of inheritable traits, including horizontal gene transfers and mutations that decrease uptake and efflux pump overexpression^13, 14, 15^. Although a number of genes and mechanisms of resistance to antibiotics are already known in *P. aeruginosa*, the complex mechanisms controlling the basal, low-level resistance to these compounds are still poorly understood^16, 17^.

The OM of *P. aeruginosa* is known to be central to its antibiotic-resistance phenotype. Its intrinsically low permeability is partly determined by inefficient OM porin proteins that provide innate resistance to several antimicrobial compounds, mainly hydrophilic ^1, 8, 10^. On the other hand, the loss of specific efflux pump mechanisms, commonly overproduced in clinical isolates, is compensated by reducing the permeability of the OM^9^. Thus, mechanisms involved in OM organization, composition and integrity interfere with the diffusion through the membrane of antimicrobial compounds, either hydrophilic or hydrophobic. Particularly the asymmetric lipid organization of the OM is the main responsible for the low permeability to lipophilic antibiotics and detergents^18^.

In *Escherichia coli*, the Mla system (MlaA-MlaBCDEF) was initially proposed to have a phospholipid import function, preventing phospholipid accumulation in the outer leaflet of the OM and thus controlling membrane-phospholipid asymmetry^19^. The core components of this ATP-binding-cassette (ABC) transport system in the inner membrane (IM) comprise the permease (MlaE), the ATPase (MlaF) and the substrate-binding protein MlaD that are highly conserved among Gram-negative bacteria^20^. The MlaA component, an integral OM protein that forms a channel adjacent to trimeric porins, is thought to selectively remove phospholipids from the outer OM leaflet and transfer them to the soluble periplasmic substrate-binding protein MlaC^21, 22^. MlaC would then transport the phospholipids across the periplasm and deliver them to MlaD for active internalization through the IM^23^. Deletion of the genes of this system is known to destabilize the OM, and bacterial strains lacking any of the Mla components are more susceptible to membrane stress agents^19, 24, 25, 26, 27, 28, 29, 30^. More recently, the retrograde transport hypothesis has been questioned and a new role for this system in anterograde phospholipid transport has been suggested^29, 31^.

The orthologous Mla system in *P. aeruginosa* is encoded by the PA4452-PA4456 operon (locus tags corresponding to PAO1) and the isolated gene PA2800 (MlaA ortholog, also known as VacJ). Proteins encoded by this gene cluster are highly similar to those encoded by operon *ttg2* (toluene tolerance genes) in *Pseudomonas putida*^32, 33^. Although it is unlikely that organic solvents themselves are substrates of this transporter, this system was initially linked to toluene tolerance in that bacterium^34^. Accordingly, components of the *P. aeruginosa* ABC transporter encoded by the PA4452-PA4456 have been named Ttg2A (MlaF), Ttg2B (MlaE), Ttg2C (MlaD), Ttg2D (MlaC) and Ttg2E (MlaB)^32^. Recent studies of mutant strains with disrupted *ttg2* or *vacJ* genes support the contribution of this ABC transport system to the intrinsic resistance of *P. aeruginosa* to antimicrobials^24, 28, 32, 35^. Yet, one of these studies has challenged the role of this system in *P. aeruginosa* as an ABC importer mediating phospholipid intermembrane trafficking^32^.

Here, we have primarily focused on *P. aeruginosa*’s Ttg2D (Ttg2D_Pae_) to further study the function of the Ttg2 system in these bacteria. We present structural and functional evidence of the role of this protein as a phospholipid transporter. Our structural analysis further enriches the existing knowledge on the structural diversity of substrate-binding proteins (SBPs) and supports current discussions on the directionality of phospholipid transport by the Mla system. In addition, based on mutational studies of the *ttg2* operon, we have validated the contribution of the Ttg2 system to the intrinsic basal resistance of *P. aeruginosa* to several antibiotic classes and other damaging compounds. Although the role of other components of this ABC transport system in multi-drug resistance has been already established for *P. aeruginosa*^24, 32^, this is the first study focusing on the soluble periplasmic SBP component, Ttg2D_Pae_. Among the components of the Ttg2 system, this SPB could be the most promising candidate for an antimicrobial intervention based on the specific blocking of this trafficking pathway.

## Results

### Ttg2D_Pae_ contains a large hydrophobic cavity that binds four acyl tails

Sequence analysis indicates that Ttg2D_Pae_ (PA4453) is the soluble periplasmic SBP component of the ABC transporter encoded by the *ttg2* operon and a member of the Pfam family MlaC (PF05494). Interestingly, the available 3D structures for the MlaC family from *Ralstonia solanacearum* (PDB entry 2QGU), *P. putida* (PDB entries 4FCZ and 5UWB) and *E. coli* (PDB entry 5UWA) were all solved in complex with a ligand in their hydrophobic pocket. Electron densities for the ligand were compatible in all cases with a phospholipid, supporting their predicted role as a phospholipid transporter. A sequence alignment shows that some of the residues thought to be involved in phospholipid binding in the *R. solanacearum* Ttg2D structure are conserved in the *P. aeruginosa* ortholog (Fig. S1). Remarkably, the electron densities for *P. putida* Ttg2D revealed the presence of two diacyl lipids in its pocket.

To investigate ligand binding at the molecular level, we determined by molecular replacement the crystallographic structure of the functional unit (aa 23-215) of Ttg2D_Pae_ at 2.53 Å resolution (PDB entry code 6HSY). The structure was refined to a final *R*_work_ and *R*_free_ of 20.9 and 24.9%, respectively, and good validation scores (Supplementary Table S1). All residues but the last three C-terminal ones (plus the C-terminal expression tag) could be modeled. Ttg2D_Pae_ adopts a mixed α+β fold with a highly twisted anti-parallel β-sheet formed by five strands and surrounded by eight α-helices. It exhibits a “decanter” shaped structure never described before for any other protein family (Fig. 1A). The structure presents a highly hydrophobic cavity between the β-sheet and the helices that spans the whole protein and has a volume of 2979 Å^3^ and a depth of ~25 Å (Fig. 1B). After the first refinement stage (AutoBuild), without any ligand added, clear density was visible inside the cavity that could correspond to four acyl chains (Fig. S2). We therefore modeled inside the cavity two PG(16:0/cy17:0) (Fig. 1A), as MALDI-TOF experiments revealed that this lipid was one of the most abundant among the different lipids found to bind to Ttg2D_Pae_ when expressed in *E. coli* (see later). Real-space correlation coefficients of 0.9 for the lipids indicate a good fit to the electron density 2*mF*_o_ - *DF*_c_. The four acyl tails are deeply inserted into the hydrophobic cavity, while the polar head groups are exposed to the solvent and make only few contacts with the protein (Fig. 1, A and C). This lack of specific recognition of the head group could explain why Ttg2D_Pae_ is able to bind different types of phospholipids. The presence of two diacyl lipids suggests that the protein could also be able to bind one tetra-acyl lipid, such as cardiolipin.

**Figure 1.**
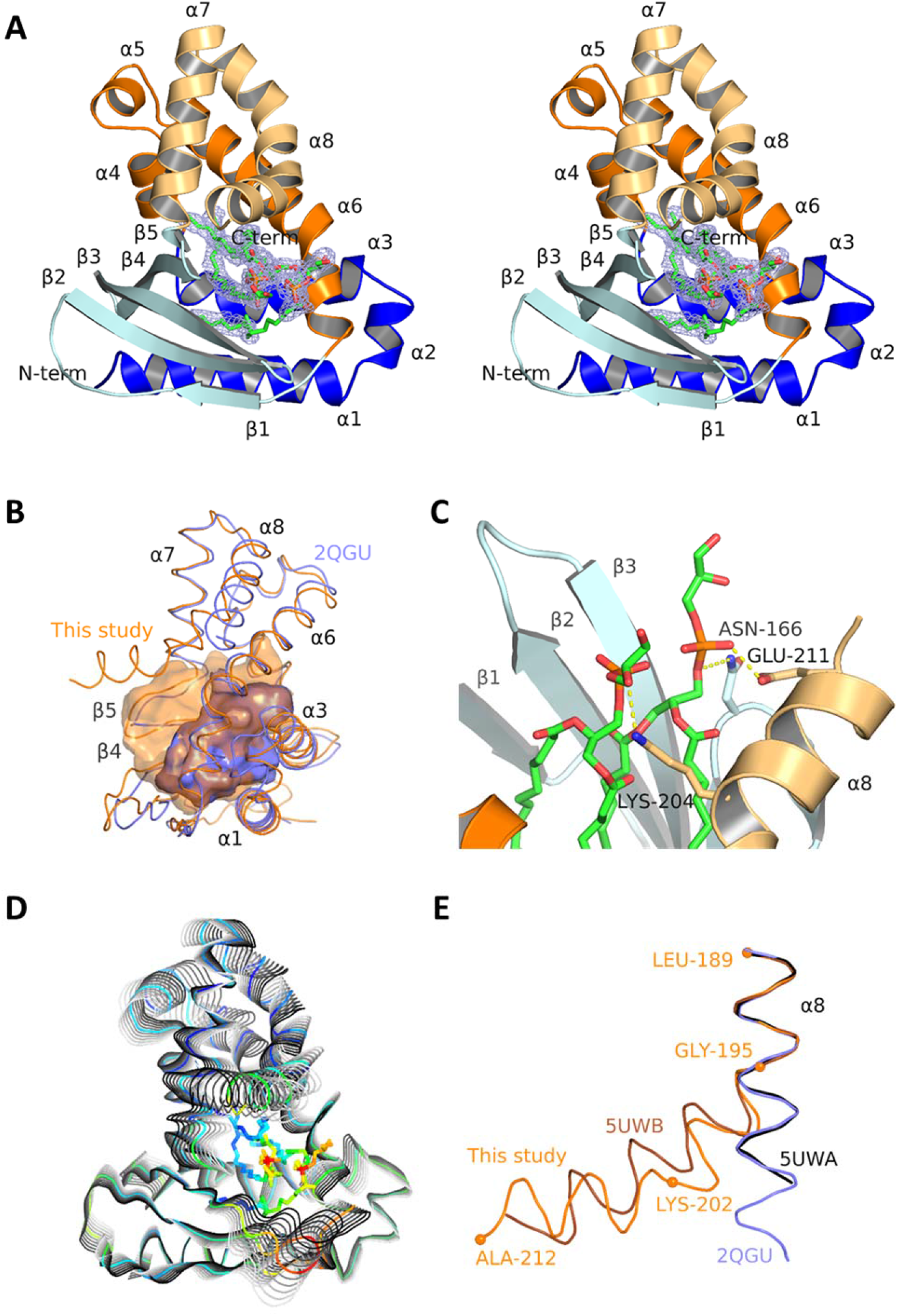
Ttg2D_Pae_ binds two phospholipids simultaneously. (A) Crystal structure of Ttg2D_Pae_ with two PG (16:0/cy17:0) bound (stereo view). The feature-enhanced electron-density map around the lipids, shown as a mesh, is contoured at 1.5σ. The cartoon representation of the protein is colored according to the CATH domains: domain 1 in blue and domain 2 in orange, dark tones for segments 1 and light tones for segments 2 in each domain (see Fig. S1). (B) Superposition of the structures of Ttg2D from *P. aeruginosa* (orange) and *R. solanacearum* (slate). The cavities of both proteins are shown as semi-transparent surfaces (side-view from the right of A). (C) Interactions between the lipid head group and the protein. (D) Protein motions along the normal mode 7. The colors represent the *B*-factors (spectrum blue to red for lowest to highest values). (E) Superposition of helix α8 of Ttg2D from *P. aeruginosa* (orange), *P. putida* (brown), *R. solanacearum* (slate) and *E. coli* (black).

To investigate the mechanism of entry and release of the two lipids in the cavity of Ttg2D_Pae_, we performed a normal mode analysis (NMA). NMA may be used to model the internal collective motions of a protein, for example upon ligand biding, generally described by a few low-frequency modes^36^. Fig. 1D shows the collective motions along mode 7, the first non-trivial mode (modes 1 to 6 account for translational and rotational motions of the protein as a whole). Rather than “en bloc” relative motions of sub-domains, all secondary structures of the protein appear to move in a concerted manner, helix α4 and the core of the β-sheet being more rigid. This breathing-like motion increases in a concerted manner the volume of the cavity and its mouth area, and may allow the lipids to enter into or exit from the cavity. Inspection of the next 10 lowest-frequency normal modes shows similar concerted motions. The normal modes can be also used to compute atomic mean-square displacements, which can be in turn related to *B*-factors^37^. The NMA-derived and the observed (crystallographic) *B*-factors are closely correlated except in regions 75-95 and 180-200, which are involved in crystal contacts, and region 105-120, where the electron density is weaker (Fig. S3). This suggests that the normal modes provide a realistic description of the protein’s flexibility.

### The 3D structure of Ttg2D_Pae_ belongs to a new SBP fold

The Ttg2D_Pae_ structure is completely different from any known, non-MlaC SBP. In general, MlaC family proteins are formed by two domains with a special segment arrangement where each domain is made by non-contiguous segments of the peptide chain (Fig. 1A and S1) in a way that resembles domain dislocation^38^. In the “decanter” shaped structure, the first domain (D1) forms the body of the decanter and adopts a Nuclear Transport Factor 2 (NTF2)-like topology (CATH Superfamily 3.10.450.50). This domain is formed by two non-contiguous sequence segments: D1S1, formed by three α-helices, and D1S2, made of five β-strands. The second domain (D2), the decanter neck, was classified as a member of a CATH superfamily (1.10.10.640) comprising only members of the MlaC family. The D2 domain is strictly all-alpha, with five helices, and it also splits in two non-contiguous sequence segments (D2S1 and D2S2) (Fig. 1A and S1).

To confirm that MlaC family proteins constitute a new fold, we have run a DALI search against the whole PDB. This returned 800 structures, all but six containing a domain of the same superfamily as D1, where the region causing the match is found. The 2OWP structure was the best non-MlaC hit. Although the reported RMSD was 2.6 Å, only 99 residues were superposed (52% of Ttg2D_Pae_) leaving out half of the protein (all residues in D2 and some in D1). In addition, the 800 DALI results were compared to a list of 501 SBP structures previously classified in different structural clusters^39^. As expected, the two lists share no common fold. We superposed the Ttg2D_Pae_ structure to a representative of each subcluster defined in the previous classification. RMSD values, number of aligned residues and structural superpositions are shown in Fig. S4. The longest match aligns 47 residues with an RMSD of 3.97 Å (2PRS chain A, a 284-residue structure member of cluster A-I), while the best RMSD is 1.41 Å with 23 residues aligned (3MQ4 chain A, a 481-residue structure member of cluster B-V). These results clearly confirm that MlaC family proteins do not belong to any previously known SBP structural cluster.

### Evolution of sequence and structural diversity of the MlaC family

Components of the Mla system are broadly conserved in Gram-negative bacteria, except for the periplasmic MlaC that notoriously shows high inter-species sequence diversity (Fig. 2A). A structural alignment of MlaC family proteins with known 3D structures (Fig. S1 and S5) reveals that, despite sequence identities ranging from 63% for the *P. putida* protein to as low as 17% for the *E. coli* one, the RMSDs of the structural alignments are very low, ranging from 1.6 to 3.1 Å (188 to 185 Cα), respectively (Supplementary Table S2). Clearly, the secondary structure elements are highly or strictly conserved among all four proteins, despite substantial amino-acid variations (Fig. S1 and S5). However, the four proteins split into two groups: *P. aeruginosa and P. putida* Ttg2D have a hydrophobic cavity of 2979-2337 Å^3^ and can bind two diacyl lipids, while the *R. solanacearum* and *E. coli* proteins have a half-size cavity of 1444-1332 Å^3^ and bind only one diacyl lipid (Supplementary Table S2). Surprisingly, although the different number of ligands had been already noticed when the structure of Ttg2D from *P. putida* was solved, cavity differences were never analyzed. Fig. 1B illustrates the cavity difference between *P. aeruginosa* and *R. solanacearum* Ttg2D. The volume differences is correlated with a different number of residues forming the cavities, from 55 down to 31 (Supplementary Table S2)However, these residues, which are spread along the whole protein sequence (Fig. S1), are largely conserved in terms of position and, in most cases, in terms of identity or similarity also, with a few substitutions such as V147/L, or V163/I or M directly affecting the volume. Some side-chain reorientations, like Y105, and small secondary structure displacements, like strands β3 and β4 or helix α6 shifted by ~2Å (Fig. S5), also modulate the volume. Taken together these changes are, nevertheless, not sufficient to explain how the cavity volume can double. The α8 helix seems to be the crucial difference between a two and a one diacyl-phospholipid cavity, not only because the helix is longer in the first case (Fig. S1), but also because it adopts a different conformation. Indeed, for the second group (*R. solanacearum* and *E. coli*, one diacyl lipid), this helix has a straight conformation (Fig. 1E), covers the α6 helix (Fig. S5) and does not participate in the cavity (Fig. 1B and S1), while in the first group (*P. aeruginosa* and *P. putida*, two diacyl lipids), the α8 helix is bent towards and over the α7 helix and greatly enlarges the cavity (with additional residues from β4 and β5 strands). This bend occurs at the conserved G195 with an angle of 40° and 64° in Ttg2D proteins from *P. aeruginosa* and *P. putida*, respectively (Fig. 1E). The helix of the first protein has an additional bend of 43° at K202. Glycine has a poor helix-forming propensity^40^ and tends to disrupt helices because of its high conformational flexibility. On the other hand, phenylalanine and glutamine have better helix-forming propensities and are found in the *R. solanacearum* and *E. coli* proteins, in which the α8 helix is straight. In addition, W196, exclusive of the pseudomonal structures, may also contribute to the influence of the α8 helix on the cavity’s volume, since its bulky hydrophobic side chain, deeply inserted into a hydrophobic pocket on the concave side of the curvature could stabilize the helix α8 bend (Fig. S5). Given our observations, we hypothesize that G195 and W196 could be crucial evolution amino-acid substitutions between one and two diacyl-phospholipid binding proteins and could be markers between the two groups. Interestingly, an alignment of 151 representative amino-acid sequences belonging to the MlaC family and identified across different Gram-negative species (Fig. 2A) revealed that G195 and W196 are conserved not only in *Pseudomonas* species but also in a group of related sequences in other non-phylogenetically related gamma-proteobacteria. In this group of proteins that hypothetically bind two diacyl phospholipids, other positions with distinct residues with respect to the whole MlaC family stand out, especially in two regions located between the central part (positions 65-83) and the C-terminal end (positions 154-198) of the protein (Fig. 2B). Side-chain orientation and hydrophobicity of some residues in these regions could be also contributing to a tighter binding of the two diacyl phospholipids inside the ligand cavity (Fig. S5). The presence of common protein sequence signatures in species that are not closely related indicates that horizontal gene transfer, mediated by recombination events between flanking conserved genes, could have contributed to MlaC family diversity.

**Figure 2.**
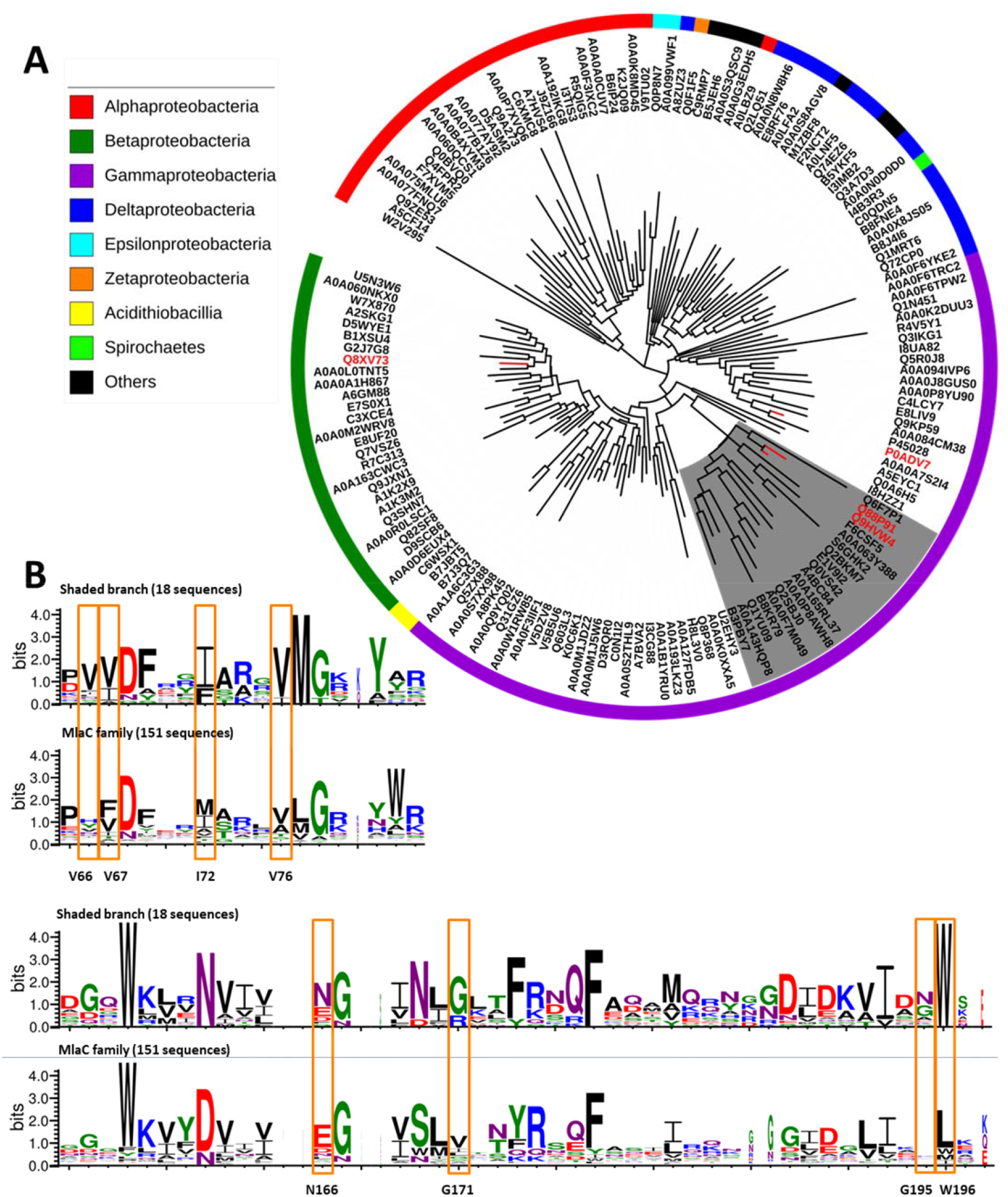
Sequence diversity among MlaC family proteins. (A) Phylogeny of 151 representative amino-acid sequences belonging to the Pfam family MlaC (PF05494) and identified across different Gram-negative species (bacterial classes are shown with different colors). The maximum likelihood tree was calculated with the LG+G+F model on MEGA 7, and visualized and annotated with iTOL. UniProt codes are used to identify each sequence, and those proteins with known 3D structure are indicated in red (P0ADV7 in *E. coli*, Q8XV73 in *R. solanacearum*, Q88P91 in *P. putida* and Q9HVW4 in *P. aeruginosa*). The branch corresponding to proteins that we predict to bind two diacyl lipids is shaded. This branch comprises sequences from different species belonging to four orders of Gamma-proteobacteria (*Pseudomonadales*, *Alteromonadales*, *Cellvibrionales* and *Oceanospirillales*). (B) Aligned sequence logos for the MlaC family in two sequence regions for the whole set of 151 representative sequences and for a sub-set of sequences corresponding to the shaded branch of the tree (18 sequences). At a given position, the height of a residue is proportional to its frequency. Residues that would distinguish the proteins binding one and two diacyl lipids are boxed.

### Ttg2D_Pae_ binds two diacyl glycerophospholipids, representing a novel phospholipid trafficking mechanism among Gram-negative bacteria

Native mass spectrometry (MS) was used to determine the biomolecules that associate noncovalently with recombinant Ttg2D_Pae_ and the stoichiometry of the interaction in a cellular environment (Fig. 3). The native mass spectrum of Ttg2D_Pae_ shows a broad charge-state distribution corresponding to the protein with multiple lipids with different masses. A major charge state with MW 24289 Da (z=9) corresponds to the delipidated recombinant protein (22819 Da) and two bound phospholipids (~1469 Da) (Fig. 3A). After isolation of selected ions (m/z 2700, z=9; m/z 2430 z=10 and m/z=2208, z=11) of intact phospholipid protein complexes (wide peak ion) and corresponding gas phase fragmentation with a transfer collision energy of 50 V, we detected the delipidated protein (m/z 2537, z=9; m/z 2283, z=10 and m/z 2075, z=11) and a family of released phospholipids (Fig.3B-C and S6). Major peaks at m/z 664.5, 704.5, 718.5 and 730.5 released from Ttg2D_Pae_ confirmed the identity of these ligands as phosphatidylethanolamines (PE) with different hydrocarbon chains (PE C30:0, PE C33:1, PE C34:1 and PE C35:2 respectively) (Fig. 3C). The dissociation experiments in the gas phase in native conditions also identified as ligands of the recombinant Ttg2D_Pae_ the major components of the bacterial membrane, phosphatidylglycerol (PG) and phosphatidylcholines (PC) (Fig. S6). In addition, released phospholipids were analyzed in positive mode denaturing conditions to ascertain their structural composition (data not shown). To characterize the largest possible number of phospholipid molecules bound by Ttg2D_Pae_, the lipid moiety of the recombinant protein was also analyzed by MS under denaturing conditions and negative ion mode (Fig. 3D). Using this method, PG C33:1 and PG C34:1 came out as most abundant, but PE C32:1, PE C33:1, PE C34:1, PG C30:0, PG C32:1, PG C32:1, PG C35:1 and PG C36:2 were also detected (Fig. 3E). The distribution of phospholipids bound to recombinant Ttg2D_Pae_ may depend on their relative abundances in *E. coli* (the recombinant protein-expression host), and it correlates well with the reported phospholipid composition of *E. coli* under comparable conditions^41, 42^. Altogether, and despite the cytoplasm of *E. coli* is clearly not the natural environment of this periplasmic protein, the total mass of the lipidated Ttg2D_Pae_ protein determined by native MS and those of the released molecules suggests that two phospholipids with different head groups could be transporter at the same time by this ABC transporter, probably also in *P. aeruginosa*.

**Figure 3.**
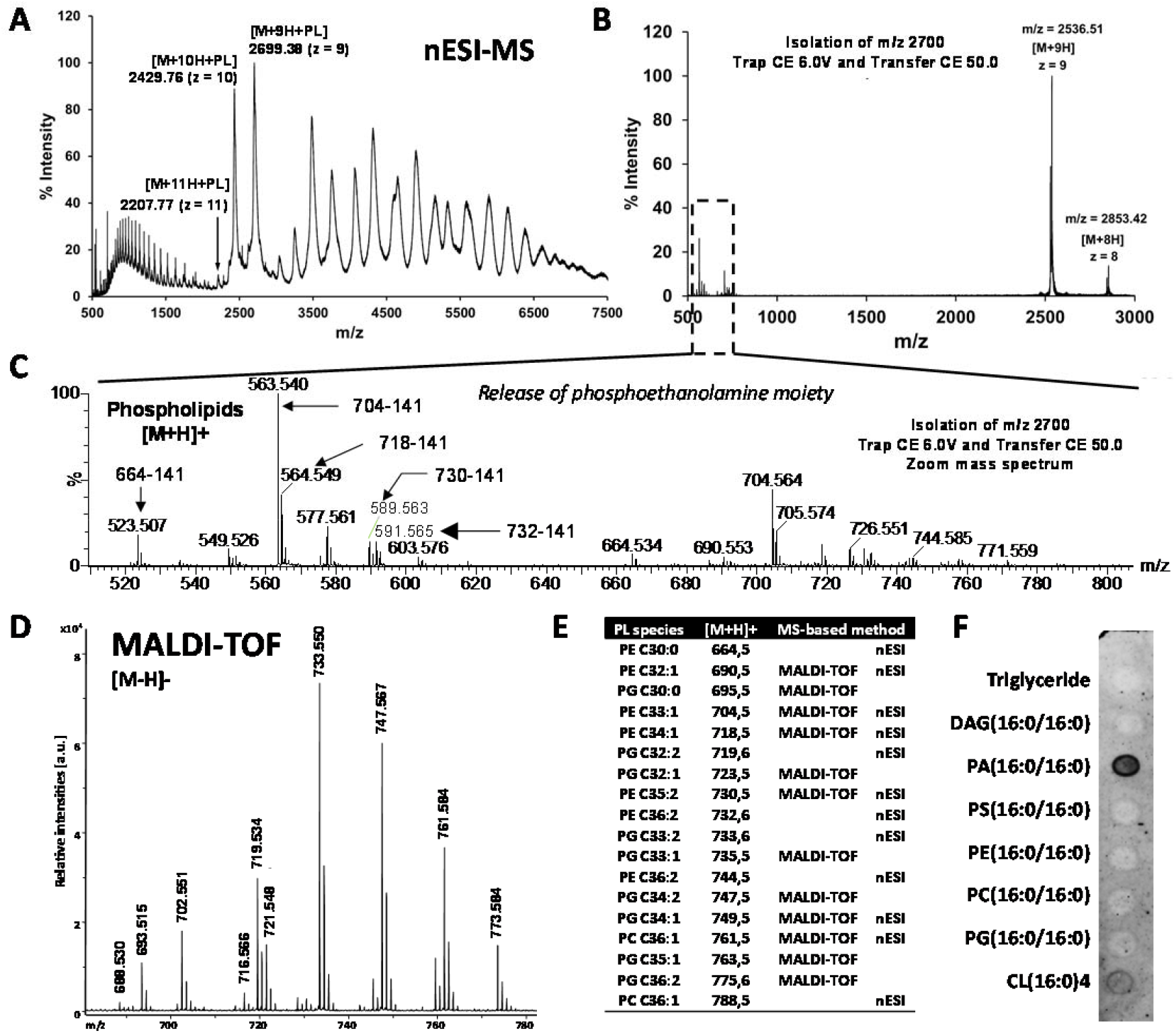
Ttg2D_Pae_ is a promiscuous phospholipid-binding protein. (A) Positive mode, non-denaturing, ESI mass spectrum of Ttg2D_Pae_ expressed in *E. coli*. The most represented species in native conditions are proteins containing at least two phospholipid (PL) molecules. (B) Mass spectrum fragmentation of isolated ion m/z = 2700 (z = 9), PLs are expected in the range 600-800 Da. (C) Zoom of the MS/MS spectrum in positive mode showing the most abundant glycerophospholipids released from Ttg2D_Pae_ under non-denaturing MS dissociation conditions. The major PLs detected show a loss of 141 Da, after MS/MS experiment, that corresponds to the release of a phosphoethanolamine (PE) moiety. (D) Negative ion mode mass spectrum under denaturing conditions of glycerophospholipids released by Ttg2D_Pae_.No other PLs were detected (data not shown). (E) Assignments of peaks in the mass spectra to different PL species (see methods). Numbers associated to each species indicate the number of carbon atoms and double bonds, respectively, in the fatty acid side chains. The most abundant ions detected by both methods correspond to two glycerophospholipid classes: phosphatidylglycerols (PG) and PE. Phosphatidylcholine (PC) species were also observed. (F) Ttg2D_Pae_ binds phospholipids in vitro. The purified delipidated protein was overlaid in an Echelon P-6002 membrane lipid strip. PA, phosphatidic acid; CL, cardiolipin; DAG, diacylglycerol; PS, phosphatidylserine.

Binding of glycerophospholipids was also demonstrated *in vitro* by testing the ability of a purified delipidated protein to bind common membrane lipids on a membrane strip (Echelon Biosciences Inc). To obtain the delipidated protein, in-column delipidation by reverse-phase liquid chromatography was used. The removal of lipids was then confirmed by MALDI–TOF and native MS analyzes (data not shown). Incubation of delipidated Ttg2D_Pae_ with the membrane lipid strip resulted in protein binding to phosphatidic acid (PA) and to a lesser extent cardiolipin (Fig. 3F). Ttg2D_Pae_ did not bind to spots on the strips containing only PE or PG. This result could support the previous suggestion that Ttg2D_Pae_ may have preference for binding two phospholipids with different head groups. It must also be taken into account that the state of the phospholipids in the spots on the membrane is far from representative of that found *in vivo*.

### The Ttg2 system provides *P. aeruginosa* with a mechanism of resistance to membrane-damaging agents

As expected, a *P. aeruginosa* Δ*ttg2D* mutant exhibited a debilitated outer membrane leading to increased susceptibility to several membrane damaging agents (Fig. 4), as demonstrated by the 1-N-phenylnaphthylamine (NPN) assay. Indeed, an enhancement in NPN uptake was observed in the mutant in the presence of the permeabilizer agents EDTA and colistin (Fig. 4, A and B). In line with this, the Δ*ttg2D* mutant is significantly more susceptible to the action of polymyxins (lipid-mediated uptake), but also of antibiotics that use both the lipid- and porin-mediated pathways to penetrate the cell, including fluoroquinolones, tetracyclines and chloramphenicol (Fig. 4C). With regards to polymyxin antibiotics, the *ttg2*D transposon insertion mutant was eight-fold more susceptible to colistin than the PAO1 wild-type, a colistin-susceptible reference strain (Table S3). In general, the mutation did not significantly affect the resistance phenotype displayed by the PAO1 strain to the beta-lactam antibiotics or aminoglycosides tested. The susceptibility phenotypes due to deletion of *ttg2D* could be fully or partially reverted by complementation with the cloned *ttg2D* gene or the full operon *ttg2* in the replicative broad-range vector pBBR1-MCS5 (Fig. 4 and Table S3), confirming the link between the gene and the phenotypes. We have also confirmed that insertional mutations in each of the other components of the *ttg2* operon (*ttg2A*, *ttg2B*, *ttg2C*) and *vacJ* (*mlaA* ortholog) lead to an increased susceptibility to antibiotics in the same way as for the Δ*ttg2D* mutant (Table S3). The Δ*ttg2D* mutant is also significantly susceptible to the toxic effect of the organic solvent xylene (Fig. 4D) and it is four-fold more susceptible to the chelating agent EDTA (MIC=0.5 mM) than the parental wild-type PAO1. However, no difference was observed between the mutant and wild-type cells in their susceptibility to SDS, obtaining for both strains a MIC value of 0.8%. Finally, disruption of the *ttg2D* gene resulted in an approximately two-fold reduction in biofilm formation and increased notoriously the activity of EDTA against *P. aeruginosa* biofilms at a subinhibitory concentration of 0.05 mM (Fig. 4E).

**Figure 4.**
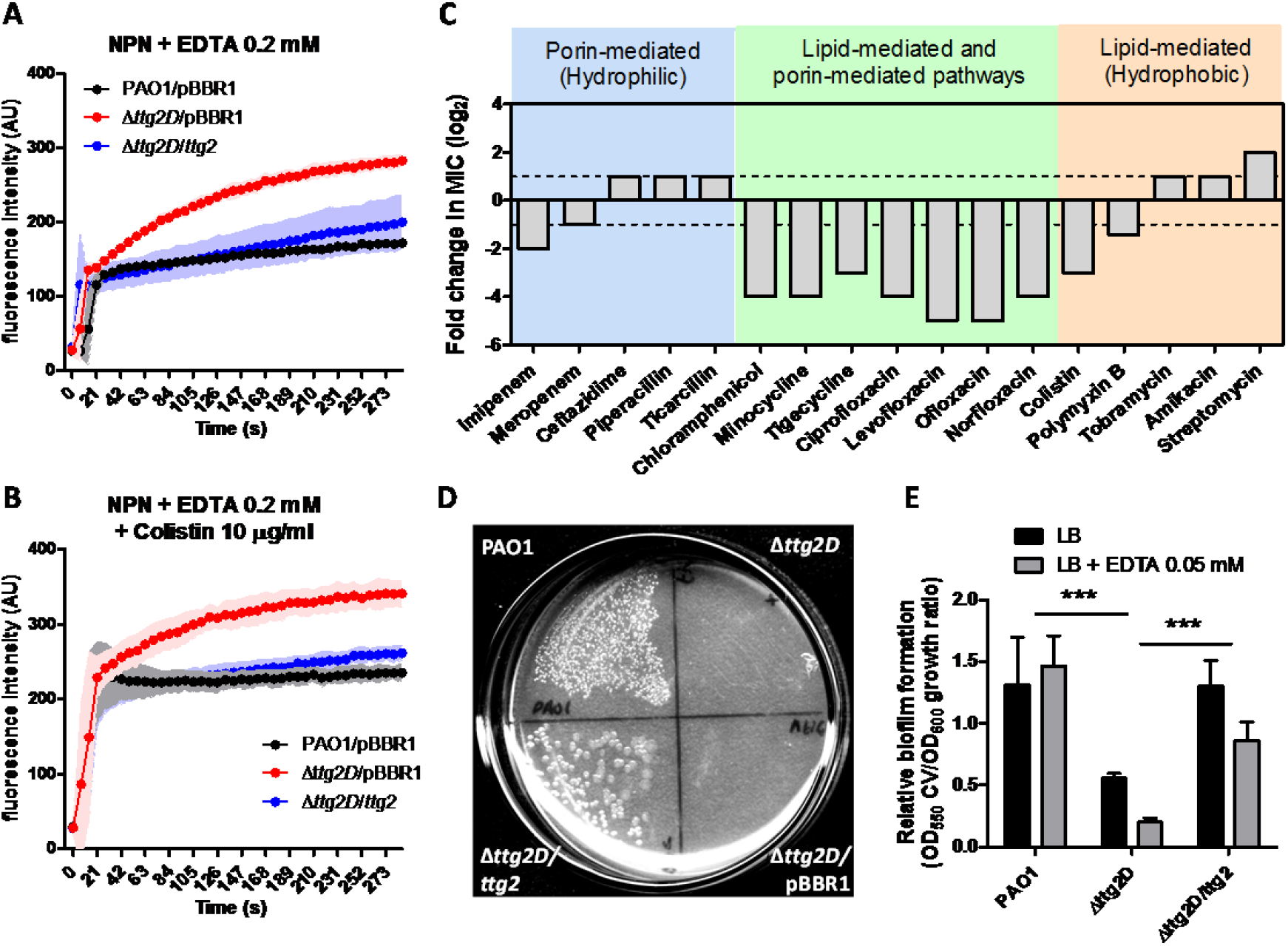
Phenotypic changes in the Δ*ttg2D P. aeruginosa* mutant denote destabilization of its outer membrane. (A and B) Ability of EDTA and colistin to permeabilize the outer membrane (NPN assay) of the native, mutant and complemented PAO1. (C) Relative change of the Δ*ttg2*D mutant MIC (minimum inhibitory concentration) for antibiotics of different classes grouped according to their cell entry mechanism. Fold changes were determined with respect to the PAO1 wild type, represented as dotted lines. (D) Growth in LBMg plates overlaid with 100% *p*-xylene. Under this condition the growth was assessed following incubation at 37°C for 24h. The image is representative of duplicate experiments. (E) Relative biofilm formation determined by crystal violet (CV) staining for Δ*ttg2D* mutant and control strains in LB medium with and without EDTA. Asterisks denote the significance of the data between groups (one-way ANOVA with Tukey’s multiple comparison test). In all panels pBBR1 indicates insertion of pBBR1MCS-5 vector alone as a control.

### The Ttg2 system is associated to *P. aeruginosa*’s intrinsic resistance to low antibiotic concentrations

The susceptibility of Ttg2-defective mutants to antibiotics was further studied in strains with different genetic backgrounds. To this end, the full *ttg2* operon was mutated in the clinical MDR *P. aeruginosa* strains C17, PAER-10821 and LESB58, which had shown different patterns of resistance to several antibiotic classes, specifically, polymyxins, fluoroquinolones and tetracyclines (Table 1). In particular, PAER-10821 and LESB58 are *P. aeruginosa* strains with low-level resistance to colistin. The generation of mutants with disrupted gene functions in MDR bacteria is troublesome because the antibiotics commonly used in the laboratory are no longer useful for selection of gene knockouts. In addition, the loci mutated in this case is involved in a general mechanism of resistance to antimicrobial agents and mutant strains are therefore expected to be generally susceptible and thus potentially lost during the selection steps. For this reason we have adapted a mutagenesis system based on the homing endonuclease I-SceI^43, 44^ to construct targeted, non-polar, unmarked gene deletions in MDR *P. aeruginosa* strains (see material and methods, text S1 and Fig. S7 for details). With this modified mutagenesis strategy we have obtained and validated unmarked deletion mutants of the selected MDR strains lacking the full *ttg2* operon (Fig. S7). Complemented strains were also obtained by transformation of mutant strains with a replicative plasmid containing the full *ttg2* operon and its expression in the complemented clones was confirmed by RT-PCR (Fig. S7). All these strains were tested for their susceptibilities to different classes of antibiotics (Table 1).

**Table 1.** Antibiotic susceptibility profile of *P. aeruginosa* MDR strains lacking the full *ttg2* operon.

| Antibiotic | MIC<sup>†</sup> in µg ml<sup>-1</sup> |  |  |  |  |  |
| --- | --- | --- | --- | --- | --- | --- |
|  | LESB58 |  | C17 |  | PAER-10821 |  |
|  | WT | Δ*ttg2* | WT | Δ*ttg2* | WT | Δ*ttg2* |
| **Polypeptides** |  |  |  |  |  |  |
| Colistin | 4 | 0.125* | 2 | 0.125* | 32 | 32 |
| **Fluoroquinolones** |  |  |  |  |  |  |
| Ciprofloxacin | 2 | 1 | 256 | 64* | 256 | 128 |
| Levofloxacin | 8 | 2* | 256 | 256 | 256 | 128 |
| Ofloxacin | 16 | 4* | >32 | >32 | >32 | >32 |
| Norfloxacin | 8 | 4 | >256 | >256 | >256 | 256 |
| **Tetracyclines** |  |  |  |  |  |  |
| Tetracycline | 16 | 8 | 32 | 16 | 32 | 8* |
| Minocycline | 32 | 8* | 16 | 8 | 32 | 8* |
| Tigecycline | 16 | 8 | 64 | 8* | 32 | 8* |
| **Chloramphenicol** |  |  |  |  |  |  |
| Chloramphenicol | 32 | 32 | 128 | 64 | 128 | 64 |
| **Sulfonamides** |  |  |  |  |  |  |
| Trimethoprim-sulphamethoxazole | 16 | 8 | >64 | >64 | >64 | 64 |
| **Aminoglycosides** |  |  |  |  |  |  |
| Tobramycin | 8 | 2* | 64 | 128 | 128 | >128 |
| Amikacin | 64 | 64 | 8 | 32* | 32 | 32 |
| Gentamicin | 32 | 16 | >128 | >128 | >128 | >128 |
| Kanamycin | >512 | >512 | 256 | 512 | 512 | 512 |
| Streptomycin | >64 | >64 | >64 | >64 | >64 | >64 |
| **Carbapenems (beta-lactam)** |  |  |  |  |  |  |
| Imipenem | 2 | 2 | 32 | 32 | 32 | 64 |
| Meropenem | 2 | 2 | 32 | 16 | 16 | 16 |
| **Cephalosporins (beta-lactam)** |  |  |  |  |  |  |
| Ceftazidime | 256 | 256 | 64 | 128 | 16 | 32 |
| **Penicillins (beta-lactam)** |  |  |  |  |  |  |
| Piperacillin | 256 | 256 | 256 | >256 | 128 | 256 |
| Piperacillin-tazobactam | 128 | 128 | 256 | >256 | 64 | 64 |
| Ticarcillin | >256 | >256 | >256 | 256 | 64 | 128 |
| Ticarcillin-clavulanic acid | >32 | >32 | >32 | >32 | >32 | >32 |

<sup>†</sup> Minimum inhibitory concentration (MIC) determined by the broth microdilution method. MICs were confirmed by two or three independent replicates and MIC differences greater than 2-fold with respect to the corresponding wild type strain were considered significant (indicated with an asterisk).

The three *ttg2* mutants were significantly more sensitive (between 4-and 64-fold) than the corresponding wild-type bacteria to colistin, fluoroquinolones, and tetracycline analogues, but not to the other antibiotic classes (Table 1). The mutant susceptibility phenotypes could be reverted by providing an intact copy of the entire PAO1 *ttg2* operon (PA4456-PA4452) in a replicative plasmid, except for colistin. The lack of complementation of the colistin susceptibility phenotype could be due to the effect of the antibiotic erythromycin (used as a selection marker for complemented strains) on the expression of global regulators that may influence colistin susceptibility^45, 46^ or to the overexpression of the *ttg2* operon components (two- to eight-fold with respect to wild type, see Fig. S7) that may also affect the distribution of phospholipids in the OM. Surprisingly, the susceptibility to amikacin significantly decreased for the C17 mutant and an opposite effect was observed for the LESB58 mutant and tobramycin, suggesting a genetic-background component in the effect of the *ttg2* mutation on the susceptibility to these antibiotics.

## Discussion

Here, we report a structural and functional study of the soluble periplasmic SBP of the Ttg2 ABC transport system in *P. aeruginosa* (Ttg2D_Pae_) that reveals new facets of this protein family and provides additional insight into the role of this pathway in *P. aeruginosa*. We have first characterized this protein at the molecular level, supporting its predicted role as a phospholipid transporter. Early studies of the ortholog Mla system in *E. coli* indicated that Mla is one of the systems responsible for the maintenance of lipid asymmetry in the Gram-negative OM, by retrograde trafficking of phospholipids from the OM to the cytoplasm through the IM^19^. The crystal structure of recombinant Ttg2D_Pae_ (Fig. 1) shows that it binds four acyl chains. Although we cannot rule out the transport of tetra-acyl species like cardiolipin, our crystallographic data, supported by MS studies, suggests the presence of two phospholipids in the crystal. Reevaluation of the Ttg2D structure from *P. putida* (PDB 4FCZ) by Ekiert et al.^26^ (PDB 5UWB) had also suggested the presence of a tetra-acyl, cardiolipin-like lipids in its hydrophobic pocket. In addition, *in vitro* binding studies of Ttg2D_Pae_ to cardiolipin reinforces the hypothesis that in *P. aeruginosa* this system also participates in cardiolipin transport (Fig. 3). This is in contrast to the orthologs from *E. coli* (PDB 5WA) and *R. solanacearum* (PDB 2QGU), which bind a single diacylglyceride. This indicates that, among Gram-negative bacteria, the ability of MlaC family proteins to transport two molecules at the same time is exclusive to some taxonomic groups. Phylogenetic and sequence analysis (Fig. 2), using G195 and W196 as signature for the larger cavity, suggest that there are other genera in addition to *Pseudomonas* where the Mla system transports two molecules simultaneously. This finding raises the question whether the evolution of this system in these species has been driven by transport efficiency (double cargo) or transport diversity (tetra-acyl in addition to diacyl phospholipids). Furthermore, are the two phospholipids translocated simultaneously by the permease Ttg2B, as it would need to be for a tetra-acyl phospholipid such as cardiolipin? The determination of the structure of additional transport components in other species will be necessary to corroborate our proposed classification and answer these questions. In addition, our results suggest that Ttg2D_Pae_ may be also able to carry PA (Fig. 3). Although not an abundant lipid constituent in bacteria, PA is an important intermediate in the biosynthesis of phospholipids, participates in phospholipid recycling and is a signaling molecule^47, 48^. However, high-affinity binding to PA could be an artifact of the method, possibly due to the way phospholipids are immobilized on hydrophobic membrane strips, since this class of phospholipid was not identified by MS.

Operon *ttg2* resembles the classic organization of an ABC importer^49^. Unlike most of the ABC exporters, ABC importers in Gram-negative bacteria require periplasmic SBPs that provide specificity and high-affinity. In addition, it is widely accepted that the direction of substrate transport of ABC transporters can be predicted on the basis of both the sequence of the nucleotide-binding component (ATPase)^49, 50^ and the transmembrane-domain fold of the permease component^51^. The close orthologs in *E. coli* and *Mycobacterium tuberculosis* of the *P. aeruginosa* ATPase Ttg2A, MlaF (60% identity) and the Mce protein, Mkl (40% identity), respectively, have sequence signatures typically found in prokaryotic ABC import cassettes^19, 49^. The remote homolog TGD3 from *Arabidopsis thaliana* is also a component of an ABC transport system (TGD) that imports phosphatidic acid to the chloroplasts through its outer and inner envelopes^52^. On the other hand, structural similarity searches for the *Acinetobacter baumannii* MlaE protein (Ttg2B, PDB 6IC4 chains G and H)^29^ (data not shown), identified as best match a structure of the human ABCA1 (PDB 5XJY), a known ATP-binding cassette phospholipid exporter^53^. In a recent study, and based on results on a Ttg2A mutant, the function of the Ttg2 system in *P. aeruginosa* was associated with the export of antibiotics such as tetracycline out of the cell^32^. Although, in our opinion, the Ttg2 system does not play a role as an antibiotic efflux mechanism, as proposed by these authors, the structural similarity of the permease to human export permeases suggests we should not rule out the possibility of anterograde, in addition to retrograde, phospholipid trafficking. One possibility would be that of a countercurrent model^54^, in which different types of phospholipids would exchange between the two membranes obeying to a gradient (Fig. 5). A countercurrent model would explain how asymmetries in lipid distribution in the two membranes might be achieved^54^. Although genetic and functional evidences have mainly suggested that the Ttg2/Mla pathway is a retrograde transport system^19, 55, 56^, recent studies in *E. coli* have shown that MlaD spontaneously transfers phospholipids to MlaC *in vitro*^31, 57^.

**Figure 5.**
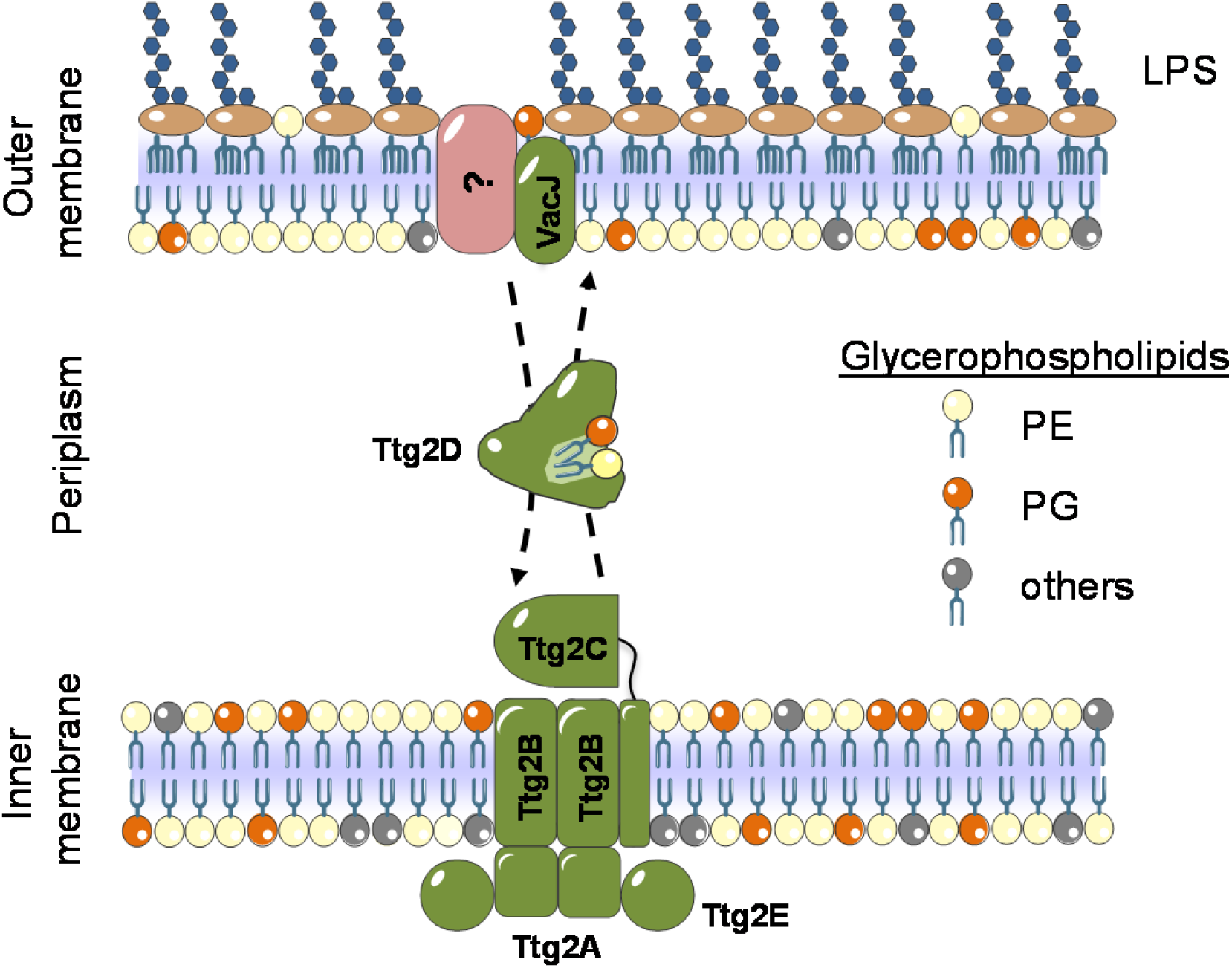
Proposed model of the Ttg2 system in *P. aeruginosa.* The soluble, periplasmic substrate binding protein Ttg2D and its orthologs in other species are thought to transport mislocalized phospholipids from the outer leaflet of the outer membrane (OM) to the inner membrane (IM) complex Ttg2ABCE across the periplasm^21^. In *P. aeruginosa*, it is not yet known if the VacJ component of the system, which delivers the lipids to Ttg2D, forms a complex with specific porins, as in *E. coli*, to extract the lipids from the membrane. In contrast to the *E. coli* ortholog (MlaC), Ttg2D_Pae_ carries two glycerophospholipids and, structure wise, could accommodate a tetra-acyl phospholipid such as cardiolipin. In addition, the protein may carry simultaneously two PL with different head groups. Considering recent studies^29, 31^, the structural signatures of the ATPase and permease models from the ortholog Mla system in *E. coli*^26^, characteristic of importer and exporter ABC cassettes, we propose, in addition to the aforementioned role in retrograde phospholipid trafficking, a second potential mode of action as an anterograde trafficking system (dashed lines) that would contribute to the maintenance of phospholipid distribution asymmetry.

MS analyses (Fig. 3) show that purified recombinant Ttg2D_Pae_ is indeed able to bind phospholipids of different chain lengths and degree of unsaturation. In addition, they indicate that this transporter may simultaneously load two phospholipids with different head groups, particularly a PG and a PE. Therefore, this system would not only control the global phospholipid content of the OM, but may also control its phospholipid composition. We have provided additional evidence, based on the NPN-uptake assay, that the Ttg2 system controls the permeability of the OM (Fig. 4). Bacterial cells tightly regulate the phospholipid composition of the OM to fortify the permeability barrier against small toxic molecules, including antibiotics. For example, anionic phospholipids like PG interact with membrane proteins and cationic antibiotics in ways that zwitterionic phospholipids like PE do not; their balance requiring a fine control^18, 58^. Indeed, the membrane’s PE content is a major factor determining the bacterial susceptibility to certain antimicrobial agents^58, 59^. In the case of positively charged antimicrobial peptides and polymyxins, it has been proposed that they promote the clustering of anionic lipids leading to phase-boundary defects that transiently breach the permeability barrier of the cell membrane^58^. In *P. aeruginosa*, an organism showing significant intrinsic resistance to certain antibiotics, the membrane PE/PG composition is approximately 60-80%/18-21%^58, 60^. Simultaneous transport of two different phospholipids across cell membranes could help control membrane charge balance and to prevent the appearance of phospholipid clusters or domains with equal charge.

Cellular studies showed that deletion of Ttg2D in *P. aeruginosa* specifically increases the susceptibility to polymyxin, fluoroquinolone, chloramphenicol and tetracycline antibiotics in the PAO1 reference strain and in three MDR clinical strains (Table 1). This mutated phenotype was observed both in the presence and absence of specific resistance mechanisms providing high-level resistance. For example, PAO1 is a relatively sensitive strain and LESB58 is a MDR strain, and both show diversity in their resistomes^61^. Thus, for the strain and antibiotic panel considered, the increase in susceptibility upon *ttg2* deletion seems to correlate with the antibiotic class rather than with the genetic background. This is in line with the physico-chemical properties of these antimicrobial compounds. Albeit positively charged, colistin is a significantly hydrophobic antibiotic that appears to gain access to the IM by permeating through the OM bilayer, while tetracyclines, chloramphenicol and quinolones use a lipid-mediated or a porin-mediated pathway depending on protonation state^62^. These antibiotic classes are classified within the same group of molecules according to their interactions with the cell permeability barriers^9^. The fact that other relatively hydrophobic antibiotics such as aminoglycosides are unaffected by the disruption of the Ttg2 system speaks in favor of the observed correlation between membrane phospholipid content and specific susceptibility to certain antibiotics^58, 59^. Another hypothesis that would explain the different impact of Ttg2 disruption on different antibiotic classes would be the possibility that components of the Ttg2 system may interact with or stabilize certain efflux pumps in *P. aeruginosa*. Indeed, the protein composition of the OM can also have a strong impact on the sensitivity of bacteria to the different antibiotic classes^62^. This aspect, however, requires further investigation.

Colistin is considered a last-resort antibiotic for the treatment of infections by several MDR Gram-negative pathogens, but its use against MDR *P. aeruginosa* is increasingly impeded by colistin resistance^63^. A variety of gene mutations are known to cause resistance to colistin by altering the OM of Gram-negative bacteria, for example, by covalent modification of the lipid A constituent of LPS as consequence of mutations in the PhoP/PhoQ two component regulatory system^64, 65^. In *P. aeruginosa*, the PhoP/PhoQ system plays a role in the induction of resistance to polymyxins in response to limiting divalent cations, as well as in virulence^66, 67^. Interestingly, this system has been recently identified as a regulator of *P. aeruginosa*’s *ttg2* operon^32^. More recently, nucleotide polymorphisms conferring resistance to polymyxins have been detected in genes of the Mla pathway in *A. baumannii*^55^. Although data on the precise mechanisms of resistance are scant and appear to be dependent on specific regulatory systems^66, 68^, the activity of the Ttg2 system on membrane phospholipid homeostasis appears to be partly responsible for the lower basal susceptibility of *P. aeruginosa* to colistin.

The proposed function of the Ttg2/Mla pathway in membrane remodeling provides a plausible explanation for the pleiotropic resistance phenotypes shown by the *ttg2* mutants in this study, including resistance to various antibiotics, chelating agents and organic solvents. In addition, these mutations increase the deleterious effect of antibiofilm agents like EDTA, a substance with known low activity against biofilms of *P. aeruginosa* PAO1^69^. Mutations in orthologous *ttg2* genes in other Gram-negative organisms have been shown to affect diverse physiological process, mainly associated with an increased OM permeability. In *E. coli*, the mutants defective in components of the Mla system rendered cells more susceptible to the lethal action of quinolones, the detergent SDS and EDTA^19, 70^. Mutants for the orthologs of the Ttg2 pathway in both *Shigella flexneri* and *Francisella novicida* resulted also in increased sensitivity to lysis by SDS^25, 71^. In addition, in *S. flexneri* this pathway appears to play a role in the intercellular spread of the bacteria between adjacent epithelial cells^25^. In fact, the Ttg2/Mla pathway has proven to be an important virulence factor in other pathogens, like *Burkholderia pseudomallei*, that need to spread into neighboring cells to infect eukaryotic tissues^72^. In *Burkholderia cepacia* complex species *mla* genes are required for swarming motility and serum resistance^28^. Furthermore, in nontypeable *Haemophilus influenzae* (NTHi), it is considered a key factor for bacterial survival in the human airway upon exposure to hydrophobic antibiotics^27^. In *S. flexneri*, *B. pseudomallei* and NTHi the role of the *mla* operon in virulence has been inferred from mutants for the gene *vacJ* (*mlaA*)^72, 73^. This gene is predicted to be part of the Ttg2 ABC transport system, since it is found in an operon with *ttg2* homologs in other bacteria^20^. While *P. aeruginosa*’s *vacJ* gene is located outside the *ttg2* operon, we have data demonstrating that strains lacking this gene share the same phenotype shown by *ttg2* mutants. In agreement with our work, it has been previously shown that in *P. aeruginosa,* VacJ plays an important role in both virulence and antibiotic susceptibility to ciprofloxacin, chloramphenicol and tetracycline^74^. In *E. coli*, this protein forms an active complex with the outer membrane proteins OmpC and OmpF^21, 23, 75^. However, in *P. aeruginosa* there are no clear orthologs to either of these porins, increasing the singular characteristics of this system in this species and suggesting potential mechanistic differences with the more studied *E. coli* transporter (Fig. 5).

## Methods

### Bacterial strains

All bacterial strains used in this study are provided in supplementary Table S4 and growth conditions in supplementary Text S1.

### Ttg2D (PA4453) structure resolution

Recombinant Ttg2D from *P. aeruginosa* was obtained in *Escherichia coli* BL21(DE3) and was purified to >99% purity. The recombinant protein obtained is tagged with a 6-histidine tail. Detailed methods for protein production, crystallization, data collection and structure refinement are available in supplementary Text S1. The data collection, processing, and refinement statistics are given in supplementary Table S1. Atomic coordinates and structure factors have been deposited in the PDB with entry code 6HSY.

### Native mass spectrometry analysis and identification of abundant phospholipids

Native MS experiments were performed using a Synapt G1-HDMS mass spectrometer (Waters, Manchester, UK) at the Mass Spectrometry Core Facility of IRB Barcelona. Prior to the analysis, samples were desalted with 100 mM ammonium acetate on centricon micro concentrator. Samples were infused by automated chip-based nanoelectrospray using a Triversa Nanomate system (Advion BioSciences, Ithaca, NY, USA) as the interface. See supplementary material (Text S1) for method details. Fragmentation of representative abundant glycerophospholipids released from Ttg2D_Pae_ was done under denaturing MS conditions (non-native). For denaturation, MS samples were directly injected to LTQ-FT Ultra mass spectrometer (Thermo Scientific, USA) using the Triversa Nanomate system. The NanoMate aspirated the samples from a 384-well plate (protein Lobind) with disposable, conductive pipette tips, and infused the samples through the nanoESI Chip (which consists of 400 nozzles in a 20×20 array) towards the mass spectrometer. Spray voltage was 1.75 kV and delivery pressure was 0.50 psi. Capillary temperature, capillary voltage and tube lens were set to 200°C, 35 V and 100 V, respectively. MS1 and MS2 spectra were acquired at 100 k resolution. Isolated ions were fragmented by CID (collision induced dissociation) with CE (collision energy) of 30 eV.

### Lipid extraction from purified recombinant protein Ttg2D and phospholipid identification

Lipid extraction from purified recombinant protein was performed according to a slightly modified version of the method described by Bligh and Dyer^76^. Briefly, 90 μl of deionized water and 750 μl of 1:2 (v/v) CHCl3:CH3OH were added to 110 μl of protein solution (∼0.5 mg ml^−1^ protein concentration) and the mixture was vortexed. After addition of 250 μl of CHCl3 and 250 μl of water, the mixture was vortexed again for 1 min and centrifuged at 1000 rpm for 5 min to give a two-phase system. The bottom phase containing the phospholipids was carefully recovered and washed with 450 μl of “authentic upper phase”. The washed bottom phase was dried in a vacuum centrifuge and dissolved in 100 μl of CHCl3 for matrix-assisted laser desorption/ionization time of flight (MALDI–TOF) MS analyses. As control, an unrelated bacterial recombinant protein, produced with the same expression system, was subjected to identical extraction protocol.

Two microliters of lipid extract were mixed with 2 μl of 9-aminoacridine (10 mg ml^−1^ dissolved in a 60:40 (v/v) isopropanol:acetonitrile solution) as MALDI matrix and 1 μl of the mixture was spotted on a ground steel plate (Bruker Daltonics, Bremen, Germany). MALDI-MS analyses were performed on an UltrafleXtreme (Bruker Daltonics) and were recorded in the reflectron negative ion mode. The ion acceleration was set to 20 kV. The spectra were processed using Flex Analysis 3.4 software (Bruker Daltonics) and they were analyzed in a mass range between m/z 450 and m/z 1,500 Da. The identification of *E. coli* phospholipids present in the sample was done according to Oursel *et al.*, 2007^41^ and Gidden *et al.*, 2009^42^, using Lipidomics Gateway (http://www.lipidmaps.org) based on the m/z values of MS spectra.

### Delipidation of purified recombinant Ttg2D

Recombinant protein, diluted 1:1 with 1% TFA, was delipidated using an HPLC system and a C18 column (Phenomenex Jupiter 5U C18 300A) in 0.1% TFA. Protein was eluted with a gradient of acetonitrile, 0.1% TFA (monitored at 214 and 280 nm) and its delipidation was checked by both MALDI-TOF and native MS analyses. Delipidated protein was lyophilized, and typically resuspended in 100 mM NaCl, 10 mM Tris-HCl (pH 8.5), to counteract the acidity of TFA, before exchanging the buffer to the desired one. Lipids bound to the column were washed out with a gradient of water-ethanol.

### Screen to identify membrane lipids binding specifically to Ttg2D

Membrane lipid strips (P-6002) were purchased from Echelon Biosciences. First, the lipid membranes were blocked in 3% (w/v) BSA in wash buffer (10 mM Tris pH 8.0, 150 mM NaCl, 1% Tween 20%) for one hour at room temperature. Second, purified delipidated recombinant Ttg2D was added in blocking buffer at a final concentration of 5 μg/ml and incubated for one hour at room temperature followed by three washes with wash buffer. As control, lipid-bound recombinant Ttg2D was used. Third, lipid membranes were incubated with a 6x-His tag polyclonal antibody HRP conjugate (MA1-21315, ThermoFisher) in blocking buffer for one hour at room temperature followed by three washes with wash buffer. Finally, the membranes were processed for enhanced chemiluminescence detection (ECL Prime Western Blotting Detection Kit) and a fluorescent image analyzer was used to detect the chemiluminescence.

### Generation of markerless *ttg2* mutants in MDR *P. aeruginosa* strains and complementations

Markerless *P. aeruginosa* mutants were constructed using a modification of the pGPI-SceI/pDAI-SceI system (Fig. S7) originally developed for bacteria of the genus *Burkholderia* and other MDR Gram-negative organisms^44, 77^. The bacterial strains and plasmids of the pGPI-SceI/pDAI-SceI system were kindly donated by Uwe Mamat (Leibniz-Center for Medicine and Biosciences, Research Center Borstel, Borstel, Germany) with permission of Miguel A. Valvano (Center for Infection and Immunity, Queen’s University, Belfast, UK). The pGPI-SceI-XCm plasmid^43^ was first modified to facilitate the generation of *ttg2* mutants in MDR *P. aeruginosa* strains. Plasmid modifications include replacement of the chloramphenicol resistance cassette by an erythromycin resistance cassette and deletion of a DNA region containing the Pc promoter found in *P. aeruginosa* class 1 integrons (Text S1 and supplementary Table S4 for details). The sequence of the new suicide plasmid vector, pGPI-SceI-XErm, is available through GenBank under the accession number KY368390. For complementation in PAO1, full *ttg2* operon or the codifying region of the *ttg2*D gene were cloned into the broad-host-range cloning vector pBBR1MCS-5 or a variant thereof containing the arabinose promoter, respectively (Table S4). For complementation experiments in MDR strains, the cloning vector pBBR1MCS-5 was first modified to confer resistance to erythromycin (see details in Text S1). Sequence for the new cloning vector, pBBR1MCS-6 is available through GenBank under the accession number KY368389. Complemented strains were obtained by transforming mutant cells with the corresponding pBBR1MCS derivative plasmid. The expression of *ttg2D* in mutant and complemented strains was verified by reverse transcription-PCR (RT-PCR) and quantitative real-time RT-PCR analysis (Text S1 and Fig. S7).

### Outer membrane permeabilization assay

Fluorometric assessment of outer membrane permeabilization was done by the 1-*N*-phenylnaphthylamine (NPN) uptake assay as described by Loh et al.^78^ with modifications (Text S1). Since *P. aeruginosa* PAO1 cells have proven to be poorly permeable to NPN^9^, either EDTA (0.2 mM) or colistin (10 μg ml^−1^) was added to cells to enhance uptake and fluorescence.

### Susceptibility to antibiotics and membrane-damaging agents

Antimicrobial susceptibility to a range of antibiotics was tested by determination of the minimum inhibitory concentration (MIC) using the broth microdilution method or Etest (Biomerieux) strips, following the Clinical and Laboratory Standards Institute (CLSI) guidelines^79, 80^ and manufacturer’s instructions, respectively (see Text S1 for details). MIC differences higher than 2-fold were considered significant changes in antibiotic susceptibility. Low-level, basal resistance to a given antibiotic was defined as that of an organism lacking acquired mechanisms of resistance to that antibiotic and displaying a MIC above the common range for the susceptible population^81^. Clinical susceptibility breakpoints against *Pseudomonas sp.* for selected antibiotics have been established by EUCAST^82^. Tolerance to organic solvents and SDS/EDTA was assessed using solvent overlaid-solid medium and MIC assays, respectively (Text S1).

### Biofilm formation

Biofilm quantification in 96-well microtiter plate by the crystal violet assay was done as previously described^83^ with modifications (supplementary Text S1).

### Bioinformatic analysis

Details are provided in the supplementary Text S1.

## Acknowledgements

This work has been supported by funding under the Seventh Research Framework Programme of the European Union (ref. HEALTH-F3-2009-223101) and the Spanish Ministry of Science, Innovation and Universities (ref. BIO2015-66674-R). The funders had no role in study design, data collection and interpretation, or the decision to submit the work for publication. We acknowledge the European Synchrotron Radiation Facility for provision of synchrotron radiation facilities and thank the staff of ID23-1 for assistance in using the beamline. Part of the mass spectrometry experiments were performed at UAB’s proteomics facility SePBioEs.

## Contributions

DY, LC, OCS, AM, MDL, MFN and MV conducted the experiments; DY, LC, OCS, MV, IG and XD designed the experiments and participated in the analysis and interpretation of experimental data; DY, LC and OCS wrote the paper; MV, IG and XD supervised research and revised the manuscript.

## Supplementary Materials

**Supplementary text S1 with methods**

**Figures S1 – S7**

**Tables S1 – S4**

**Supplementary references**

