## Supplementary materials for "Evolution toward maximum transport capacity of the Ttg2 ABC system in *Pseudomonas aeruginosa*"

**Figures S1 – S7**

**Tables S1 – S4**

**Supplementary text S1 with methods**

**Supplementary references**

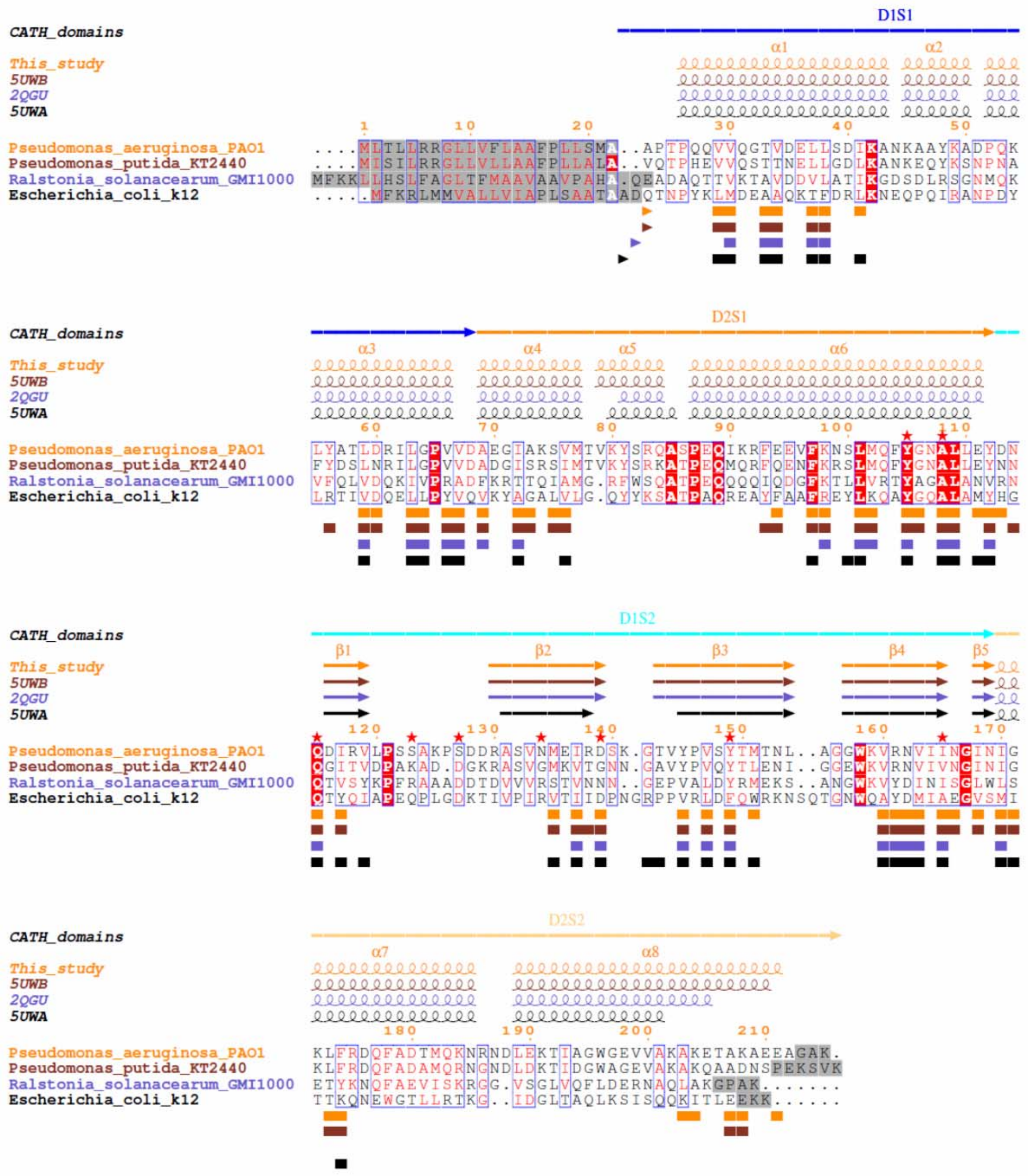


**Figure S1.** Multiple-sequence alignment of MlaC family proteins with known 3D structure based on a structural alignment. CATH domains 1 (D1) and 2 (D2), each with segments 1 (S1) and 2 (S2), are shown above the sequences together with their secondary structures elements. Identical residues are highlighted in red, similar residues are in red and similar regions are boxed in blue Residues highlighted in grey are missing in the PDB entries. Sequence numbers correspond to *P. aeruginosa* protein (Ttg2D$_{Pae}$). Red stars indicate residues annotated in the binding site of MlaC from *R. solanacearum* (2QGU). Below the sequences, triangles indicate the first amino acid of the mature protein (after cleavage of the signal peptides) and squares, the residues forming the cavities.

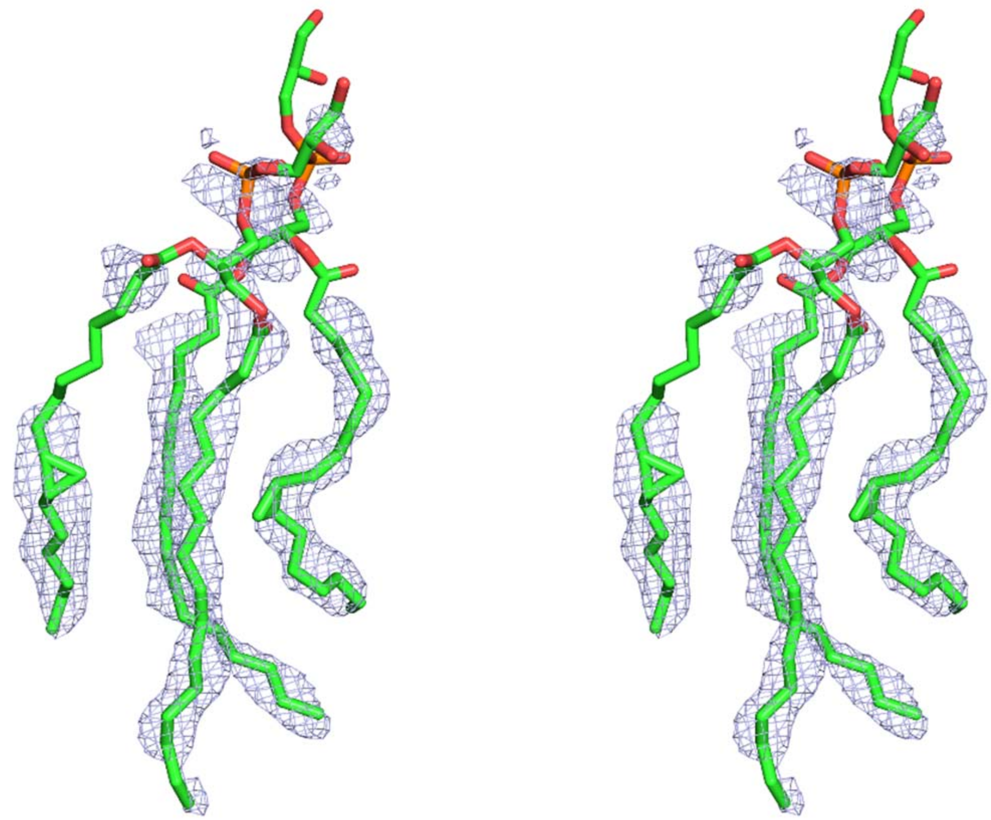

**Figure S2.** Unbiased $2mF_o$ - $DF_c$ electron density map from AutoBuild (without any ligand added) for the refined phospholipids, contoured at 1σ (stereo view).

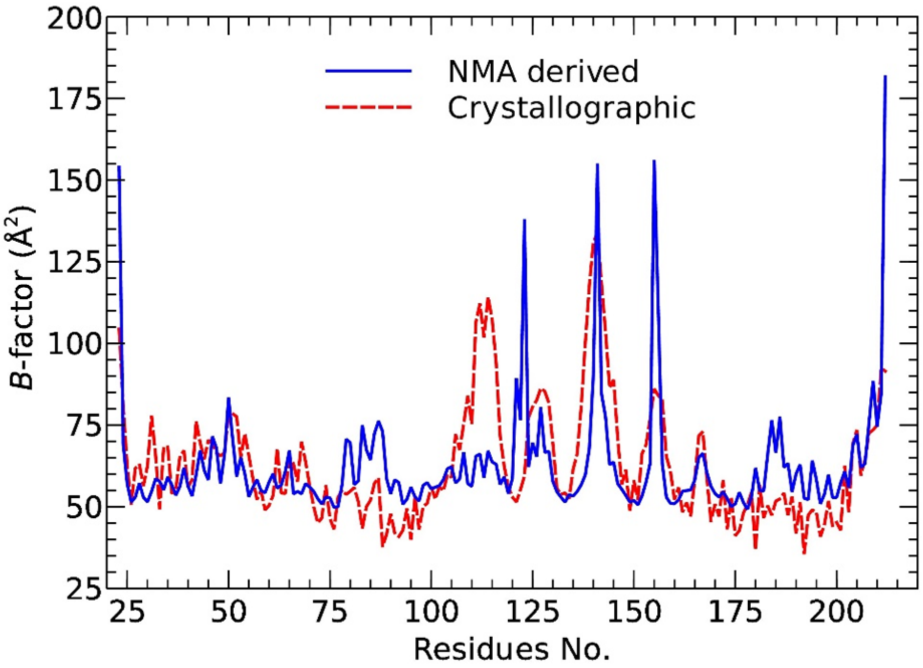


**Figure S3.** Correlation between normal-mode-derived and crystallographic mean-square displacement parameters (Cα *B*-factors).

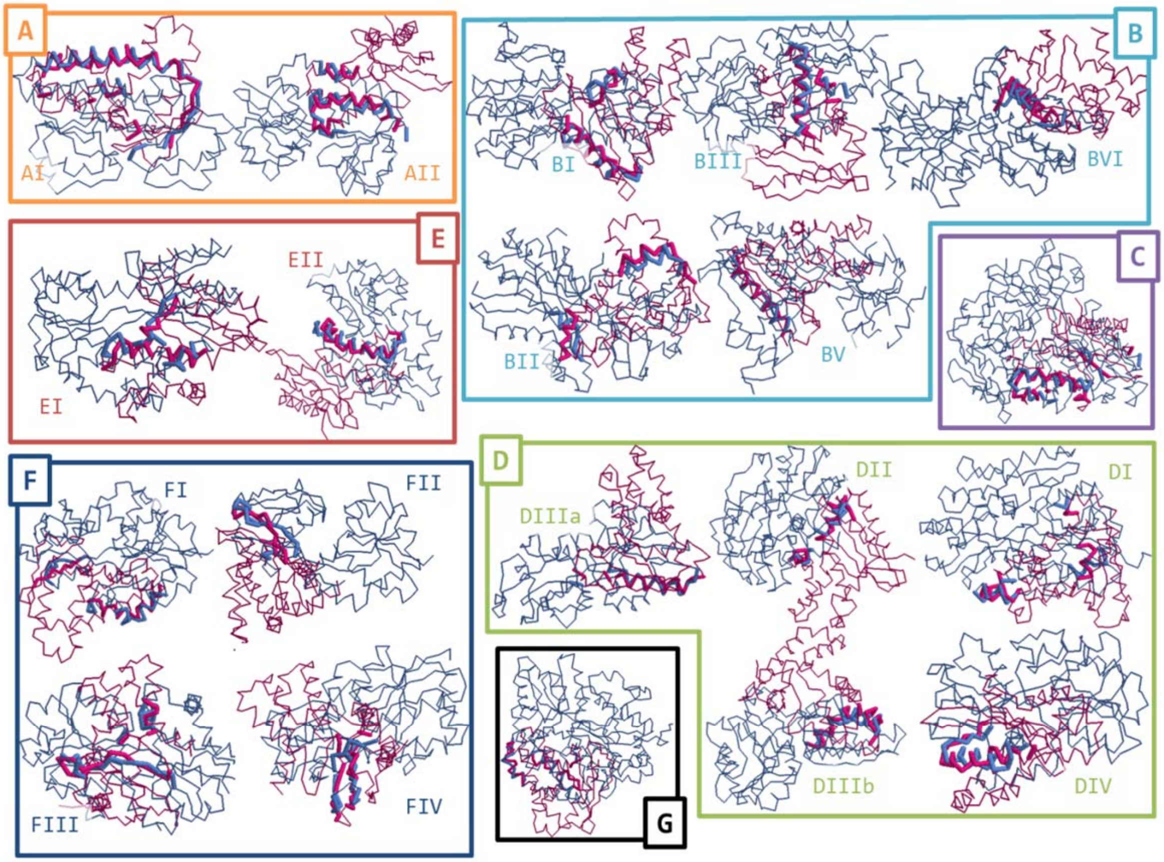


| SBP PDB_Cluster | Aligned residues | RMSD Å | Fig. panel | SBP PDB_Cluster | Aligned residues | RMSD Å | Fig. panel |
|---|---|---|---|---|---|---|---|
| 2prs_A-I | 47 | 3,97 | A | 4exl_D-IIIa | 22 | 2,29 | D |
| **2x4l_A-II** | **37** | **3,77** | **A** | **3cg1_D-IIIb** | **25** | **3,26** | **D** |
| 3s99_B-I | 30 | 2,87 | B | 1o7t_D-IV | 25 | 3,6 | D |
| **3om0_B-II** | **24** | **3,94** | **B** | **2cex_E-I** | **26** | **3,53** | **E** |
| 3sg0_B-III | 29 | 3,88 | B | 2qpq_E-II | 27 | 3,17 | E |
| **1jdn_B-IV** | **24** | **2,5** | **B** | **2x7p_F-I** | **28** | **3,73** | **F** |
| 3mq4_B-V | 23 | 1,41 | B | 4ntl_F-II | 19 | 3,73 | F |
| **2wok-C** | **40** | **3,76** | **C** | **1r9l_F-III** | **25** | **3,42** | **F** |
| 2z8d_D-I | 24 | 3,55 | D | 1us4_F-IV | 28 | 3,84 | F |
| **2qry_D-II** | **21** | **3,51** | **D** | **1y3p_G-A** | **31** | **3,75** | **G** |

**Figure S4:** Superposition of Ttg2D$_{Pae}$ (purple structures) on a representative of each subcluster (gray structures) defined in the SBP classification by G. H. Scheepers[1]. Superposed residues are shown in thick Cα-trace. The table lists the RMSD values and the number of aligned residues for each structural alignment.

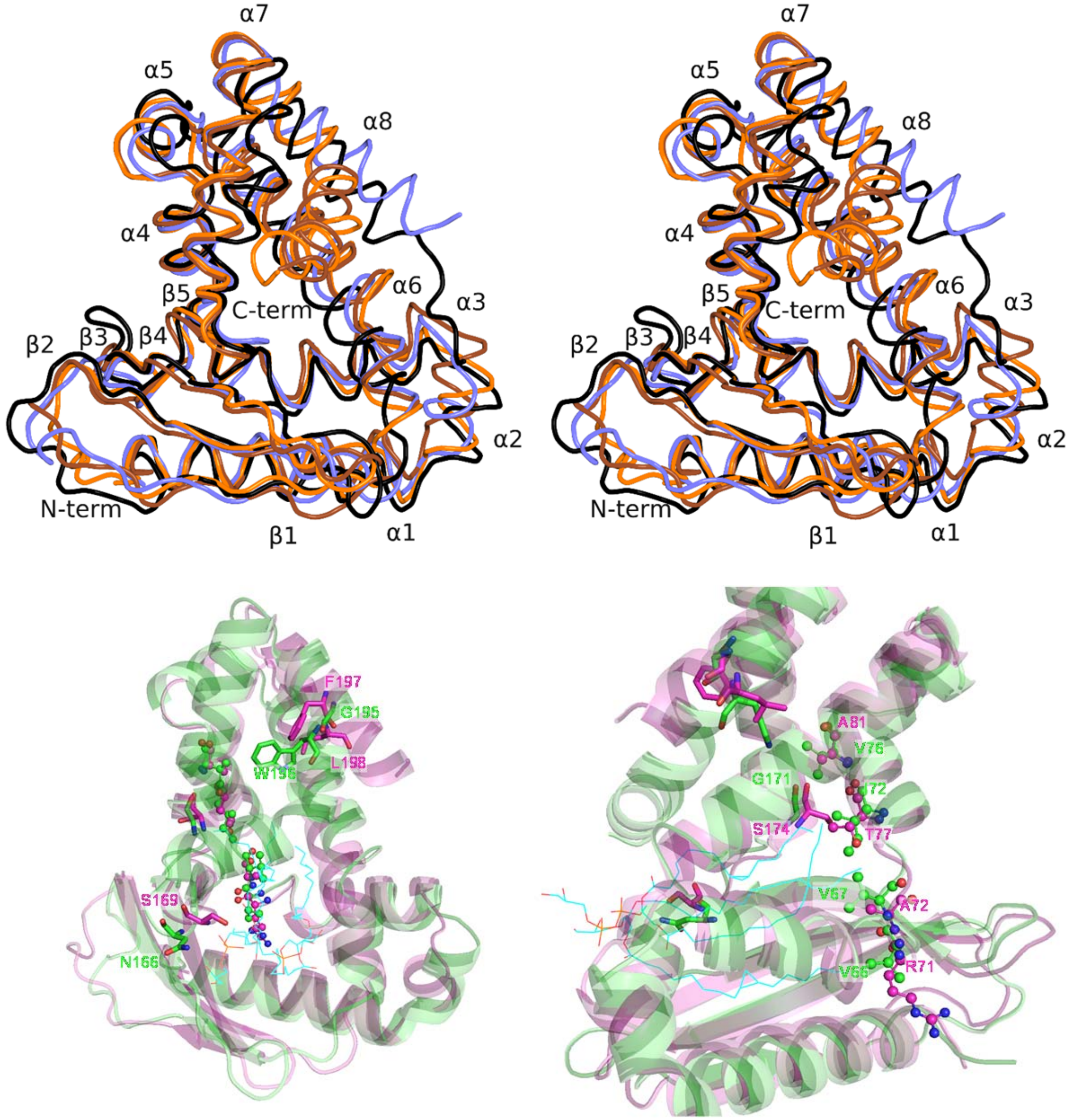


**Figure S5.** Superposition of the structure of Ttg2D from *P. aeruginosa* with known orthologous protein structures. Upper panels show superposition of Ttg2D crystal structures from *P. aeruginosa* (this study, PDB entry 6HSY), *P. putida* (5UWB), *R. solanacearum* (2QGU) and *E. coli* (5UWA) (stereo view). Same colour coding as in Fig. S1. Lower panels show superposition of the structures of Ttg2D from *P. aeruginosa* (green) and *R. solanacearum* (pink) in two different views. Residues that distinguish the group of proteins that we predict would bind two diacyl lipids are indicated with residue letter and number. Residues in *region* 65-83 are represented as *ball-and-stick and* residues in *region* 154-198 as sticks.

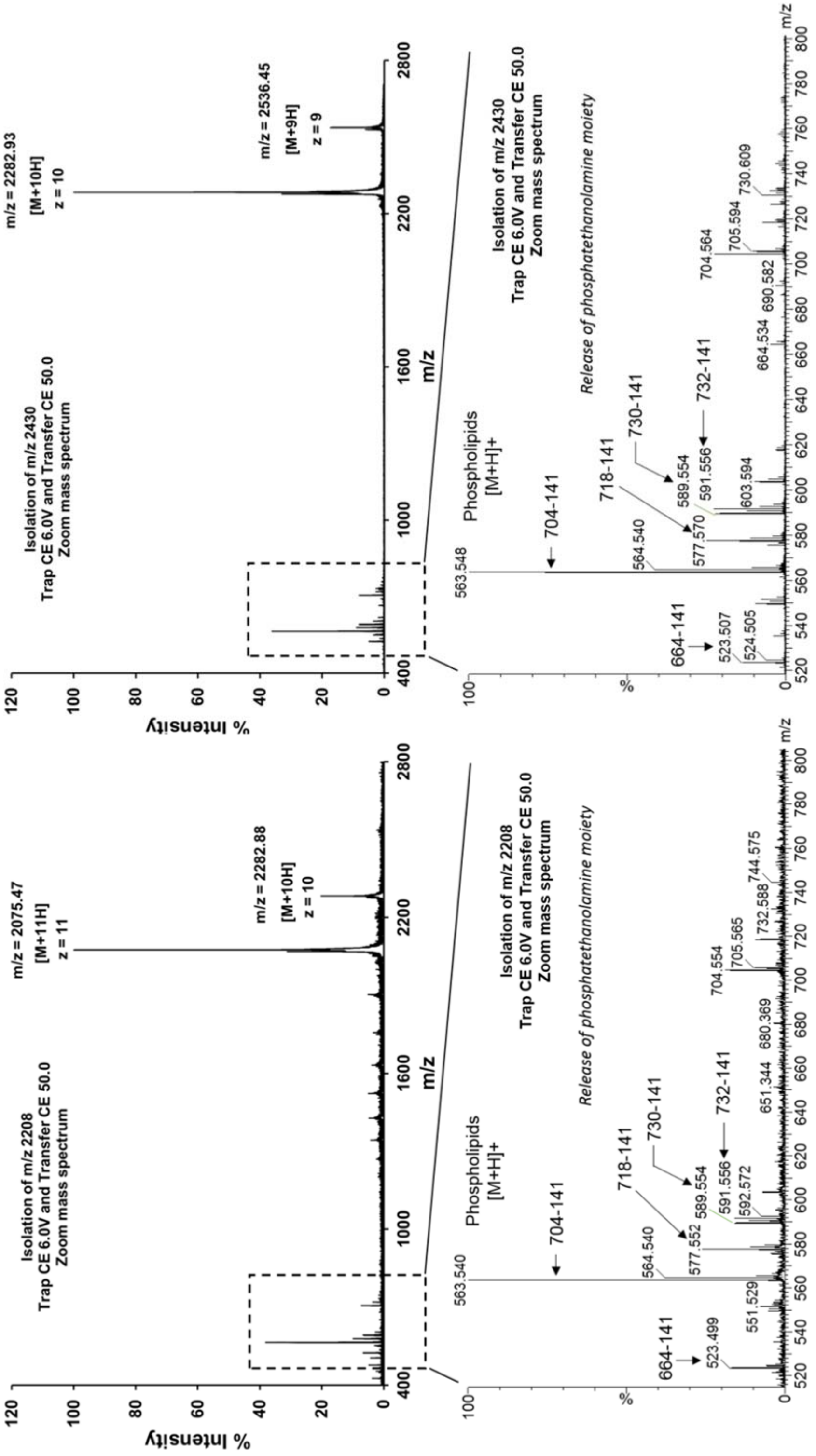


**Figure S6.** Mass spectrum fragmentation of isolated ions m/z = 2208 (z = 11) and m/z = 2430 (z = 10), and zoom of the MS/MS spectrum in positive mode showing the most abundant glycerophospholipids released from Ttg2D$_{Pae}$ under non-denaturing MS dissociation conditions. The major PLs detected show a loss of 141 Da, after MS/MS experiment, that corresponds to the release of a phosphoethanolamine (PE) moiety.

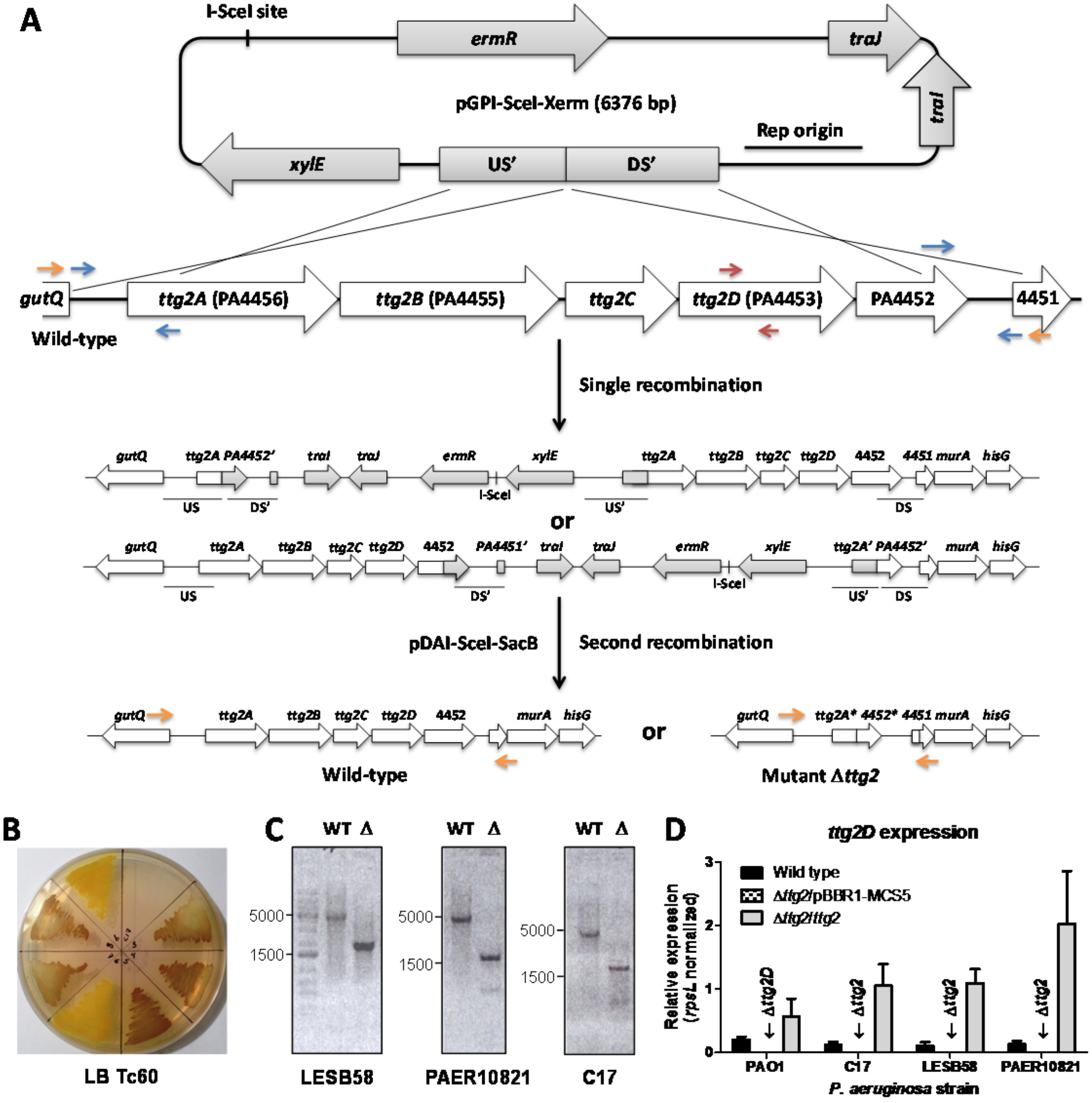


**Figure S7:** Generation of *ttg2* mutants in MDR *P. aeruginosa* strains using a modified pGPI-SceI/pDAI-SceI-SacB system. (A) Schematic representation of the construction of the *ttg2* knockout mutant. See supplementary methods for details. Primer pairs for amplification of the upstream (US) and downstream (DS) fragments for recombination events are represented as blue arrows. The second recombination event generates the desired mutant or results in reversion to wild type (WT). (B) Screening for the spontaneous resolution of cointegrates in *P. aeruginosa* strain C17 in the presence of pyrocatechol. In the presence of this compound, colonies expressing 2,3-catechol-dioxygenase encoded by *xylE* turn yellow, while *P. aeruginosa* *xylE* negative colonies turn dark brown. The picture shows a selective agar plate containing representative colonies after spraying with pyrocatechol. (C) Screening by PCR to confirm the deletion using external primers represented as orange arrows in panel A. Expected amplicon sizes in the WT and in the mutant (Δ) are 4755 bp and 1849 bp respectively. (D) Real-time PCR (qPCR) of cDNA amplified from WT and *ttg2* mutants and complemented strains using *ttg2D* internal primers (represented as red arrows in panel A). Expression levels were normalized to the *rpsL* levels.

**Table S1. Data collection and model refinement statistics**

| | Ttg2D – two PG(16:0/cy17:0) |
|---|---|
| Data collection | |
| Wavelength (Å) | 0.9802 |
| Space group | $P3_221$ |
| Unit cell parameters (Å) | $a = b = 124.64$, $c = 38.06$ |
| Resolution range (Å) | 62.32 - 2.53 (2.64 - 2.53)$^{\#}$ |
| No. of reflections | |
| Total | 71,596 (6,246) |
| Unique | 11,489 (1,349) |
| Completeness (%) | 99.6 (98.0) |
| Average multiplicity | 6.2 (4.6) |
| $\langle I/\sigma(I) \rangle$ | 8.2 (2.0) |
| $R_{meas}$ (%)$^{\dagger}$ | 13.5 (79.5) |
| $R_{pim}$ (%)$^{\ddagger}$ | 5.3 (35.2) |
| $CC_{1/2}$ (%)$^{\S}$ | 99.6 (71.7) |
| $B$-factor from Wilson plot (Å$^2$) | 43.0 |
| Model refinement | |
| No. of reflections used | 11462 |
| $R_{work}$ / $R_{free}$ (%)$^{\P}$ | 20.9 / 24.9 |
| No. of non-H atoms (all / protein / ligands / water) | 1675 / 1527 / 127 / 21 |
| No. of protein residues / chain per a.u. | 190 / 1 |
| RMS deviations | |
| Bond lengths (Å) | 0.003 |
| Bond angles (º) | 0.47 |
| Average B-factors (Å$^2$, all / protein / ligands / water) | 68.0 / 66.9 / 83.2 / 61.0 |
| Molprobity scores | |
| Overall score, %ile | 0.78, 100$^{th}$ |
| Clashscore, %ile | 0.31, 100$^{th}$ |
| Poor rotamers (%) | 1.21 |
| Ramachandran Outliers (%) | 0 |
| Ramachandran Favoured (%) | 97.87 |
| PDB code | 6HSY |

$^{\#}$Values in parentheses are for the highest resolution shell.

$^{\dagger}R_{meas} = \Sigma_{\mathbf{h}} (n_{\mathbf{h}}/(n_{\mathbf{h}}-1))^{1/2} \Sigma_i |I_{\mathbf{h},i} - \langle I_{\mathbf{h}} \rangle| / \Sigma_{\mathbf{h}}\Sigma_i I_{\mathbf{h},i}$ , where $n_{\mathbf{h}}$ is the number of observations of reflection $\mathbf{h}(hkl)$, $I_{\mathbf{h},i}$ the $i$th measurement of its intensity and $\langle I_{\mathbf{h}} \rangle$ the average of all $I_{\mathbf{h},i}$.

$^{\ddagger}R_{p.i.m.} = \Sigma_{\mathbf{h}} (1/(n_{\mathbf{h}}-1))^{1/2} \Sigma_i |I_{\mathbf{h},i} - \langle I_{\mathbf{h}} \rangle| / \Sigma_{\mathbf{h}}\Sigma_i I_{\mathbf{h},i}$.

$^{\S}$Correlation coefficient between intensities from random half-data sets.

$^{\P}\Sigma_{\mathbf{h}} ||F_o| - |F_c|| / \Sigma_{\mathbf{h}} |F_o|$, where $|F_o|$ and $|F_c|$ are observed and calculated structure factor amplitudes, respectively. $R_{free}$ was calculated using 5% of the reflections, which were not used for refinement.

**Table S2. Structural alignment parameters and cavity volumes of Ttg2D proteins**

| Species (strain) | PDB code | RMSD in Å (No. Cα) ‡ | Identity‡ | Similarity‡ | Volume† in Å$^3$ | No. cavity's aa/atoms† | Ref. |
|---|---|---|---|---|---|---|---|
| *Pseudomonas aeruginosa* (PAO1) | 6HSY | 0.0 (190) | 100% | 100% | 2979 | 55/185 | This study |
| *Pseudomonas putida* (KT2440) | 5UWB | 1.6 (188) | 63% | 78% | 2510 | 52/166 | [2] |
| | | | | | 2337 | 49/153 | |
| *Ralstonia solanacearum* (GMI1000) | 2QGU | 2.3 (180) | 25% | 48% | 1332 | 31/101 | N/A |
| *Escherichia coli* (K12) | 5UWA | 3.1 (185) | 17% | 39% | 1444 | 43/118 | [2] |
| | | | | | 1369 | 43/111 | |

‡Structural alignment parameters obtained from POSA web server for the PDB sequences.

†Molecular-surface volume (without H atoms) and the number of amino acid residues (aa) and atoms in the cavity for protein chain (chains A and B in *P. putida* and *E. coli*).

**Table S3. Antibiotic susceptibility profile of *P. aeruginosa* mutants of the Ttg2/VacJ system and complemented strains for Δ*ttg2*D.**

| **Antibiotic** | **MIC[†] in µg ml$^{-1}$** | | | | | | | |
|---|---|---|---|---|---|---|---|---|
| | **PAO1** | **Δ*ttg2*D** | **Δ*ttg2*C** | **Δ*ttg2*B** | **Δ*ttg2*A** | **Δ*vacJ*** | **Δ*ttg2D* /pBBR 1-MCS-5-*ttg2*** | **Δ*ttg2D*/ pBBR1 -pBAD-*ttg2*D** |
| Polypeptides | | | | | | | | |
| Colistin | 0.5 | 0.0625* | 0.0625* | 0.0156* | 0.0312* | 0.0625* | 0.5 | 0.25 |
| Polymyxin B | 2 | 0.75* | 1 | 1 | 1 | 1 | 4 | ND |
| Fluoroquinolones | | | | | | | | |
| Ciprofloxacin | 1 | 0.0625* | 0.0625* | 0.0312* | 0.0312* | 0.0625* | 0.125* | 0.125* |
| Levofloxacin | 4 | 0.125* | 0.125* | 0.0625* | 0.0625* | 0.125* | 0.5* | 0.25* |
| Ofloxacin | 8 | 0.25* | 0.25* | 0.25* | 0.5* | 0.5* | 1* | 0.5* |
| Norfloxacin | 4 | 0.25* | 0.25* | 0.25* | 0.25* | 0.25* | 0.5* | 0.5* |
| Tetracyclines | | | | | | | | |
| Minocycline | 32 | 2* | 2* | 2* | 2* | 2* | 8* | 8* |
| Tigecycline | 32 | 4* | 4* | 4* | 4* | 4* | 16 | 16 |
| Chloramphenicol | | | | | | | | |
| Chloramphenicol | >256 | 32* | 32* | 16* | 32* | 32* | 64* | 64* |
| Aminoglycosides | | | | | | | | |
| Tobramycin | 0,25 | 0.5 | 0.5 | 0.5 | 0.5 | 0.5 | 2 | 2 |
| Amikacin | 2 | 4 | 4 | 2 | 4 | 4 | 4 | 8 |
| Streptomycin | 8 | 32* | 64* | 32* | 32* | 32* | 64 | 64 |
| Carbapenems (beta-lactam) | | | | | | | | |
| Imipenem | 4 | 1* | 1* | 2 | 1* | 1* | 1* | 1* |
| Meropenem | 1 | 0.5 | 0.5 | 1 | 0.5 | 0.5 | 1 | 0.25* |
| Cephalosporins (beta-lactam) | | | | | | | | |
| Ceftazidime | 1 | 2 | 4* | 2 | 2 | 2 | 2 | 2 |
| Penicillins (beta-lactam) | | | | | | | | |
| Piperacillin | 2 | 4* | 8* | 8* | 16* | 8* | 4 | 8* |
| Ticarcillin | 16 | 32 | 32 | 32 | 32 | 32 | 16 | 32 |

[†]Minimum inhibitory concentration (MIC) determined by the broth microdilution method except for polymyxin B that was determined by Etest (ND not determined). MICs were confirmed by two or three independent replicates and MIC differences greater than 2-fold with respect to the wild type strain were considered significant (indicated with an asterisk). Transposon mutants of *P. aeruginosa* PAO1 and complemented strains are described in supplementary Table S4.

**Table S4:** Bacterial strains, plasmids and oligonucleotides used to study the role of the *ttg2* operon in *Pseudomonas aeruginosa*.

| Strain name | Genotype | Description | Reference |
|---|---|---|---|
| *Pseudomonas aeruginosa* | | | |
| MPAO1‡ | Wild type | Subline of PAO1. Strain lacking a transposon insertion. PAO1 is the standard laboratory and genetic reference strain. | [3] |
| PAO1Δ*ttg2D* | *ttg2D*-D04::IS*phoA*/hah (PW8497)† | Ttg2D (PA4453) mutant; Tet$^{r}$ | [3, 4] |
| PAO1Δ*ttg2C* | *ttg2C*-A12::IS*phoA*/hah (PW8498)† | Ttg2C (PA4454) mutant; Tet$^{r}$ | [3, 4] |
| PAO1Δ*ttg2B* | *ttg2B*-A07::IS*lacZ*/hah (PW8500)† | Ttg2B (PA4455) mutant; Tet$^{r}$ | [3, 4] |
| PAO1Δ*ttg2A* | *ttg2A*-B04::IS*phoA*/hah (PW8503)† | Ttg2A (PA4456) mutant; Tet$^{r}$ | [3, 4] |
| PAO1Δ*vacJ* | *vacJ*-H08::IS*phoA*/hah (PW5688)† | VacJ (PA2800) mutant; Tet$^{r}$ | [3, 4] |
| LESB58 § | Wild type | A highly virulent epidemic strain (LES) first identified in the Liverpool CF clinic center, β-lactam resistant. | [5, 6] |
| LESB58Δ*ttg2* | Δ*ttg2ABCDE* | LESB58 carrying a deletion in *ttg2* operon. | This study |
| C17 ¶ | Wild type | Clinical isolate from rectal swabs. | This study |
| C17Δ*ttg2* | Δ*ttg2ABCDE* | C17 carrying a deletion in *ttg2* operon. | This study |
| PAER-10821 ¶ | Wild type | Human clinical isolate. | This study |
| PAER-10821 Δ*ttg2* | Δ*ttg2ABCDE* | PAR10821 carrying a deletion in *ttg2* operon. | This study |
| *Escherichia coli* | | | |
| DH5α | F$^{-}$ Φ80*lacZ*ΔM15 Δ(*lacZYA-argF*) U169 *recA1 endA1 hsdR17*($r_K^-$ $m_K^+$) *phoA supE44 thi-1 gyrA96 relA1*λ$^{-}$ | For cloning purposes. | [7] |
| HY327 | Δ(*lac pro*) *argE*(*Am*) *recA56 rif*$^{R}$ *nalA* λ *pir* | Expresses the λ Pir protein required for cloning and propagation of plasmids with the R6K origin of replication. | [8] |
| BL21(DE3) | F$^{–}$ *ompT gal dcm lon hsdS*$_{B}$($r_B^- m_B^-$) λ(DE3 [*lacI lacUV5-T7p07 ind1 sam7 nin5*]) [*malB*$^{+}$]$_{K\text{-}12}$(λ$^{S}$) | Host for recombinant protein expression from plasmids containing T7 promoter. | Novagen |
| Plasmid | | Description | Reference |
| pGPI-SceI-XCm | | Mobilizable suicide vector; carries the R6Kγ origin of replication, the I-SceI recognition site and a *xylE* reporter gene, Cm$^{r}$, Tp$^{r}$ | [9] |
| pGPI-SceI-XErm | | Modified pGPI-SceI-XCm vector, Erm$^{r}$, Tp$^{r}$ | This study |
| pΔ*ttg2*-US' | | pGPI-SceI-XErm with a 633-bp *Xba*I/*Xho*I insert of PAO1 containing the flanking region upstream of *ttg2* | This study |
| pΔ*ttg2*-US'DS' | | pΔ*ttg2*-US with a 818-bp *Xho*I/*EcoR*I insert of PAO1 containing the flanking region downstream of *ttg2* | This study |

| | | | |
|---|---|---|---|
| pRK2013 | | RK2-derived helper plasmid carrying the *tra* and *mob* genes for mobilization of plasmids containing *oriT*, Kan$^r$ | [10] |
| pDAI-SceI-SacB | | Mobilizable broad host range plasmid; carries the gene for the I-SceI homing endonuclease and the *sacB* gene, Tet$^r$ | [9, 11] |
| pBBR1MCS-5 | | Broad-host-range cloning vector used for complementation, low copy, Gm$^r$ | [12] |
| pBBR1MCS-5-*ttg*2 | | pBBR1MCS-5 with the *ttg2* operon (PA4456-4452) inserted between sites *Xba*I and *Kpn*I (opposite orientation of lacZ promoter transcription), Gm$^r$ | This study |
| pBBR1MCS-6 | | Modified pBBR1MCS-5 vector, Erm$^r$ | This study |
| pBBR1MCS-6-*ttg*2 | | pBBR1MCS-6 with the *ttg2* operon (PA4456-4452) inserted between sites *Xba*I and *Kpn*I, Erm$^r$ | This study |
| pBAD18-Cm | | Expression vector containing the arabinose pBAD promoter and *araC*, Cm$^r$ | [13] |
| pBBR1-pBAD-Gm | | pBBR1MCS-5 containing the arabinose pBAD promoter and *araC* from pBAD18-Cm, Gm$^r$ | This study |
| pBBR1-pBAD-*ttg2D* | | pBBR1-pBAD-Gm with the *ttg2D* (PA4453) CDS inserted between sites *Nhe*I and *Hind*III, Gm$^r$ | This study |
| pET28b | | Bacterial expression vector with T7-lacO promoter, hexa His tag (Nterm and Cterm) with Thrombin cleavage (N terminal on backbone), Kan$^r$ | Novagen |
| Primer name | Sequence 5' to 3' | Description | Reference |
| US'-ttg2-U | GACGGAATTCTGGGCGGAATGGATGAAATC | Upstream forward primer to create pΔ*ttg2*-US', *EcoR*I | This study |
| US'-ttg2-L | TATGCTAGCTCAGCCGCAGCAACGTGGTC | Upstream reverse primer to create pΔ*ttg2*-US', *Nhe*I | This study |
| DS'-ttg2-U | CCAGCTAGCGCAGCCTTCTGGAGATCCTG | Downstream forward primer to create pΔ*ttg2*-US'DS', *Nhe*I | This study |
| DS'-ttg2-L | GACAGATCTCCGCGGATATGCAGGTCGAC | Downstream reverse primer to create pΔ*ttg2*-US'DS', *Bgl*II | This study |
| Ext-ttg2-U | GAGGTCGGGGCCAGGTTCAGTG | Forward primer outside the deleted region for mutant verification | This study |
| Ext-ttg2-L | CCTTAGCCTTGATGTAGCCGCCTTC | Reverse primer outside the deleted region for mutant verification | This study |
| Int-ttg2D-U | CAAGGCCGATCCGCAAAAGCTC | Forward primer for RT-PCR (amplicon size of 215 bp) | This study |
| Int-ttg2D-L | GCACGCGGATGTCCTGGTTGTC | Reverse primer for RT-PCR (amplicon size of 215 bp) | This study |
| ttg2compF | GTGTCTAGAGCGGAATGGATGAAATCG | Forward primer for cloning operon *ttg2* (PA4456-PA4452) into pBBR1MCS-5, *Xba*I | This study |
| ttg2compR | ATAGGTACCTTAACGTCTTCGGCCTGC | Reverse primer for cloning operon *ttg2* (PA4456-PA4452) into pBBR1MCS-5, *Kpn*I | This study |
| Erm5'-PstI | AGACTGCAGGAAACGTAAAAGAAGTTATG | Forward primer to amplify erythromycin resistance cassette, used to create pGPI-SceI-XErm, *Pst*I | This study |
| Erm3'-PstI | GAACTGCAGTACAAATTCCCCGTAGGC | Reverse primer to amplify erythromycin resistance cassette, used to create pGPI-SceI-XErm, *Pst*I | This study |

| | | | |
|---|---|---|---|
| Erm5'-KpnI | AGAGGTACCGAAACGTAAAAGAAGTTATG | Forward primer to amplify erythromycin resistance cassette, used to create pBBR1MCS-6, *Kpn*I | This study |
| Erm3'-BglII | GAGAGATCTTACAAATTCCCCGTAGGC | Reverse primer to amplify erythromycin resistance cassette, used to create pBBR1MCS-6, *Bgl*II | This study |
| pBAD18-Up | CCCACTAGTATGTCGGCGATATAG | Forward primer for cloning pBAD promoter and *araC* from pBAD18-Cm into pBBR1MCS-5, *Spe*I | This study |
| pBAD18-Lw | ATGCTCGAGGGAAATGTTGAATAC | Reverse primer for cloning pBAD promoter and *araC* from pBAD18-Cm into pBBR1MCS-5, *Xho*I | This study |
| ttg2DcompF | CCAGCTAGCGAGGTTTTCTTCCATGCTG | Forward primer for cloning *ttg2D* (CDS and RBS) into pBBR1-pBAD-Gm, *Nhe*I | This study |
| ttg2DcompR | TTCAAGCTTGCTGGCCTGGCTCATTTCG | Reverse primer for cloning *ttg2D* (CDS and RBS) into pBBR1-pBAD-Gm, *Hind*III | This study |
| PA4453-Up | ACACCATGGCTCCGACCCCGCAACAG | Forward primer for cloning *ttg2D* (mature protein) into pET28b, *Nco*I | This study |
| PA4453-Lw | CCTAAGCTTTTTCGCCCCGGCCTCTTC | Reverse primer for cloning *ttg2D* (mature protein) into pET28b, *Hind*III | This study |

† Genotype for UW mutants referenced in the following way: gene name-well name as the allele number::Transposon name. tetA, tetracycline-resistance gene. kan, kanamycin-resistance gene. All mutants contained either an ISlacZ/hah or an ISphoA/hah transposon insertion. Between parenthesis strain name at the UW mutant library [http://www.gs.washington.edu/labs/manoil/libraryindex.htm].

‡ MPAO1 (PAO1 for short) was received from the distributor of the PAO1 mutant library of the University of Washington, Seattle [3].

Note: Further information on UW mutants can be found at http://www.gs.washington.edu/labs/manoil/libraryindex.htm. The correct insertion of the transposon into the mutant strains was confirmed in the recent sequence-verified collection of UW mutants [4] and for mutants showing differential phenotypes the transposon location was also confirmed by colony PCR following the protocol and primers recommended by the University of Washington Genome Science Center.

§ The Liverpool epidemic strain (LES) B58, known as LESB58, was kindly donated by Dr. Roger C. Levesque, IBIS, Université Laval, Québec (Canada).

¶ *P. aeruginosa* C17 was isolated from an ICU patient at Hospital Clinic, Barcelona (Spain), upon screening of surveillance rectal swabs in August 2007. *P. aeruginosa* MDR strain PAR-10821 was also isolated at the Hospital Clinic in 2012.

**Text S1: Supplementary methods.**

**Bacterial growth conditions.** Unless stated otherwise, strains were routinely cultured on Luria-Bertani broth (LB) agar plates, or to exponential phase ($OD_{550}$ of 1.0), or up to late exponential phase ($OD_{550}$ of 2.7 to 3.0) in LB at 37°C with shaking at 250 rpm. When necessary, antibiotics were added at final concentrations of 500 µg $ml^{-1}$ for erythromycin, 10 µg $ml^{-1}$ for gentamicin, 17 µg $ml^{-1}$ for tetracycline, 5 µg $ml^{-1}$ for norfloxacin, or 50 µg $ml^{-1}$ for kanamycin for *Escherichia coli* or 1000 µg $ml^{-1}$ for erythromycin, 40 µg $ml^{-1}$ for gentamicin, or 60 µg $ml^{-1}$ for tetracycline for *Pseudomonas aeruginosa*.

**Ttg2D recombinant protein expression and purification.** Ttg2D (PA4453) coding sequence from PAO1 strain was cloned from residues 23 (N-terminal) to 215 (C-terminal) followed by a His6 tag into a modified pET28b expression vector (primers listed in Table S4). Expression was done in *E. coli* BL21(DE3) in LB broth by inducing with 1 mM IPTG for 3 h. Cells were disrupted by sonication in lysis buffer (0.1% Triton X-100, 4 µg $ml^{-1}$ Lysozyme, 8 µg $ml^{-1}$ DNase and 2 mM $MgCl_2$), supplemented with a tablet of Protease Inhibitor Cocktail (Roche) per 10 ml of buffer. Recombinant protein in soluble fraction was purified by two chromatographic methods, using an ÄKTA Purifier (GE Healthcare Life sciences). Protein was first purified by metal ion affinity chromatography, using a HisTrap HP 5 ml column (GE Healthcare Life sciences) with the following buffers:binding buffer: 5 mM imidazole, 0.5 M NaCl and 20 mM Tris-HCl (pH 7.9); washing buffer: 40 mM imidazole, 0.33 M NaCl and 13.3 mM Tris-HCl (pH 7.9) and elution buffer: 125 mM imidazole, 62.5 mM NaCl and 2.5 mM Tris-HCl (pH 7.9). Eluted protein was immediately dialyzed against 50 mM $Na_2HPO_4$ (pH 7.0) to remove the imidazole and then subjected to size-exclusion chromatography using a HiLoad 26/600 Superdex 75 pg column (GE Healthcare Life sciences). Protein concentration was determined by Bradford method and protein purity was evaluated by SDS-PAGE. Purification gave a very good yield, obtaining 99 mg of protein with a purity >99%.

**Crystallisation, data collection and structure refinement.** Purified Ttg2D was concentrated to 18.8 mg $ml^{-1}$ in 100 mM NaCl, 10 mM Tris-HCl (pH 7.5) and sitting-drop crystallization trials were performed at 20° C using commercial screens. Best diffracting crystals were obtained from drops of 200 nL of protein solution plus 200 nL of reservoir solution consisting of 0.17 M ammonium sulfate, 25.5% PEG 4K and 15% glycerol. Harvested crystals were directly flashed cooled in liquid nitrogen. X-ray diffraction data were collected at 100 K on the beamline ID23-1 at the European Synchrotron Radiation Facility (Grenoble, France) [14]. These data were indexed, integrated, scaled and merged using iMOSFLM [15] and *AIMLESS* [16]. Ttg2D structure was solved with Phaser [17] using a poly-alanine model built with MODELLER [18] from the homologous protein of *Ralstonia solanacearum* (PDB code 2QGU, 25% sequence identity). The structure was automatically re-built with one run of AutoBuild, followed by iterative cycles of restrained refinement with Phenix.refine [19], model building/solvent addition with *Coot* [20] and validation with MolProbity [21]. Geometry restraint information for the phospholipid PG(16:0/cy17:0) was generated from its SMILES description with eLBOW and the semi-empirical quantum mechanical method AM1 [22]. Feature-enhanced map [23] was used to build the lipids as the $2mF_o$ - $DF_c$ electron density was weak in this region. Crystallographic data and refinement statistics are reported in Table S1. Cavities and normal modes were analyzed with the web servers CASTp [24] and ElNémo [25], respectively. The structural alignment was determined with POSA [26] and rendered with ESPript 3 [27] (the PDB sequences were completed to correspond to the UniProt ones). Structural figures were prepared with PyMOL (The PyMOL Molecular Graphics System, Version 1.8 Schrödinger, LLC).

**Ion mobility spectrometry-mass spectrometry (IM-MS) analyses.** IM-MS experiments were performed using a Synapt G1-HDMS mass spectrometer (Waters, Manchester, UK). All samples were in 100 mM ammonium acetate and were infused by automated chip-based nanoelectrospray using a Triversa Nanomate system (Advion BioSciences, Ithaca, NY, USA) as the interface. The ionization was performed in positive mode using a spray voltage and a gas pressure of 1.75 kV and 0.5 psi, respectively. The source pumping speed in the backing region (6.70 mbar) of the mass spectrometer was reduced to achieve optimal transmission of non-covalent complexes. Cone voltage, extraction cone and source temperature were set to 40 V, 3 V and 40ºC, respectively. Trap and transfer collision energies were set to 6 V and 4 V, respectively, for TOF MS analysis. After ion isolation (m/z 2700, z=9; m/z 2430 z=10 and m/z=2208, z=11), fragmentation was performed by CID in the transfer region by applying a 50 V collision energy (TOF MS/MS analysis). The pressure in the Trap and Transfer T-Wave regions were $5.93 \cdot 10^{-2}$ mbar of Ar and the pressure in the IMS T-Wave was $3.99 \cdot 10^{-4}$ mbar of $N_2$. Trap gas flow was 1.5 ml/sec. The bias voltage for entering in the T-wave cell was 15 V. The instrument was calibrated over the m/z range 300-8000 Da using a solution of cesium iodide. MassLynx version 4.1 SCN 704 and Drift scope version 2.4 software were used for data processing. The experimental optimized parameters are listed in detail in the table.

| NanoESI + V Resolution mode | | DC potentials (V) | |
|---|---|---|---|
| *m/z* range | 300-8000 | Trap Collision Energy | 6 |
| Spray voltage (kV) | 1.75 | Transfer Collision Energy | 4 |
| Gas pressure (psi) | 0.5 | Trap DC Entrance | 5 |
| Source Temperature ($^{o}$C) | 40 | Trap DC Bias | 15 |
| Sampling Cone (V) | 40 | Trap DC Exit | 5 |
| Extraction Cone (V) | 3 | IMS DC Entrance | 5 |
| Desolvation temperature ($^{o}$C) | 250 | IMS DC Exit | 2 |
| Cone Gas Flow (l/hr) | 10 | Transfer DC Entrance | 2 |
| Desolvation Gas Flow (l/hr) | 300 | Transfer DC Exit | 2 |
| Trap Gas Flow (ml/min) | 8 | | |
| IMS Gas Flow (ml/min) | 25 | | |
| Backing region (mbar) | 6.70 | | |
| Trap and Transfer T-Wave section pressure (mbar of Ar) | $5.93 \cdot 10^{-2}$ | | |

**Generation of *ttg2* mutants in MDR *P. aeruginosa* strains using the pGPI-SceI/pDAI-SceI-SacB system.** This mutagenesis method is based on the I-SceI homing endonuclease system, which relies on two independent crossover events to integrate first a deletion plasmid with a I-SceI recognition site into the genome of the recipient and then resolve the co-integrate structure by a second homologous recombination event in the presence of the I-SceI endonuclease provided in trans on a replicative plasmid [11, 28]. One of these mutagenesis systems relies on an improved suicide vector that contains an I-SceI restriction site and the *xylE* reporter gene (pGPI-SceI-XCm), and a replicative but unstable plasmid that encodes the I-SceI endonuclease and the counterselectable marker SacB (pDAI-SceI-SacB) [9]. To successfully achieve genetic manipulations in MDR *P. aeruginosa* strains, we further modified pGPI-SceI-XCm by introducing an erythromycin resistance determinant replacing the chloramphenicol resistance cassette. To make this decision, the MIC for erythromycin was previously analyzed in our *P. aeruginosa* strains using the microdilution technique and found to be at a level of 256 µg $ml^{-1}$. The erythromycin resistance (*erm*) gene from plasmid pNZerm [29] was PCR amplified using primers Erm5'-PstI and Erm3'-PstI (Table S4) and with pNZerm DNA as a template. The resulting amplicon (1026 bp) was digested with *Pst*I and cloned into *Pst*I-digested pGPI-SceI-XCm to create pGPI-SceI-XErm. Previously, the region between the *Sac*I sites of plasmid pGPI-SceI-XCm (359 bp) was deleted by digestion with this restriction enzyme and ligation. This region contains the Pc promoter found in class 1 integrons (promoter for the trimethoprim resistance gene *dhfrIIb* in pGPI-SceI vectors) and this could cause unwanted integration of the suicide vector into the *P. aeruginosa* chromosome of some strains. Class 1 integrons have been detected with high prevalence in *P. aeruginosa* [30].

The mutagenesis plasmid for the *ttg2* operon deletion (from PA4452 to PA4456 in PAO1) was constructed by PCR amplification of DNA fragments flanking this gene cluster from PAO1 strain, which were cloned into pGPI-SceI-XErm (see Table S4 for primer details). The upstream fragment (633 bp) was amplified using primers US'-ttg2-U and US'-ttg2-L. The downstream fragment (818 bp) was amplified using primers DS'-ttg2-U and DS'-ttg2-L. The upstream fragment was digested with *EcoR*I and *Nhe*I, the downstream fragment was digested with *Nhe*I and *Bgl*II, and both fragments were inserted in two successive cloning steps into pGPI-SceI-XErm to create pΔ*ttg2*-US'DS'. The successful construction of the mutagenesis plasmid was verified by DNA sequence analysis of the inserts. All deletion plasmids were generated and maintained in *E. coli* SY327.

The mutagenic plasmid pΔ*ttg2*-US'DS' was mobilized into *P. aeruginosa* C17, PAER-10821 and LESB58 by triparental mating [28] using *E. coli* DH5α carrying the plasmid pRK2013 as a helper strain. Modifications of the method include that the recipient *P. aeruginosa* strains were incubated at 42°C overnight before conjugation because certain strains presumably contain restriction systems that could severely restrict foreign DNA. Erythromycin at 1000 μg $ml^{-1}$ was used to select for cointegrants (single-crossover clones) in these *P. aeruginosa* strains, and 5 μg $ml^{-1}$ norfloxacin to counter-select against the *E. coli* helper and donor strains. To distinguish true cointegrants from colonies that spontaneously became resistant to erythromycin, streaks of exconjugants were sprayed with 0.45M pyrocatechol since in the presence of this compound colonies expressing 2,3-catechol-dioxygenase encoded by *xylE* turned bright yellow [31]. For the final mutagenesis stage, pDAI-SceI-SacB was mobilized into *P. aeruginosa* cointegrants, and exconjugants were selected on LB agar plates containing 60 μg $ml^{-1}$ tetracycline and 5 μg $ml^{-1}$ norfloxacin for PAER-10821 and LESB58 derivatives. Tetracycline-resistant colonies, appearing after 48 hours, were screened by PCR and sequencing to confirm the deletion using the primers Ext-ttg2-U and Ext-ttg2-L (Table S4) that anneal to sequences outside the deleted region. Detection of deletion mutants cured from the plasmid pDAI-SceI-SacB was achieved by growing *P. aeruginosa* on LB plates without salt and supplemented with 5% (wt/vol) sucrose and then screening the resulting colonies for loss of tetracycline resistance. To obtain a *ttg2* mutant in the *P. aeruginosa* C17 strain, the derivative cointegrates have to be resolved by spontaneous recombination of the allele pair since resolution via the I-SceI endonuclease provided on the plasmid pDAI-SceI-SacB did not work. Failure during the second recombination event for C17 strain (resolution always restores the parental allele) was probably due to the mutation in the *ttg2* operon making the cells more susceptible to tetracycline [32], the antibiotic resistance marker to select for cells carrying plasmid pDAI-SceI-SacB. In this case, resolution of cointegrates was achieved by plating several thousand colonies on LB plates without selection. For this, the single-crossover clones were serially subcultured in LB without selection for two consecutive days and then diluted up to $10^{-8}$ prior to spread over the plates. Plates were sprayed with 0.45M pyrocatechol to screen for *xylE* negative cells (no yellow coloration). The screening was facilitated because *P. aeruginosa xylE* negative colonies turned dark brown after spraying with pyrocatechol (Fig. S7). Selected colonies were screened by PCR to confirm the deletion using the primers Ext-ttg2-U and Ext-ttg2-L.

**Vectors for complementation**. For complementation purposes of the PAO1 derivative mutant strains, the broad-host-range cloning vector pBBR1MCS-5 [12] was used with the oligonucleotide primers described in supplementary Table S4 for each cloning strategy. For complementation of the MDR *P. aeruginosa* strains C17, PAER-10821 and LESB58, we previously modified pBBR1MCS-5 by introducing an erythromycin resistance determinant replacing the gentamicin resistance cassette. The *erm* gene from plasmid pNZerm [29] was PCR amplified using primers Erm5'-KpnI and Erm3'-BglII (Table S4) and with pNZerm DNA as a template. The resulting amplicon (1026 bp) was digested with *Kpn*I and *Bgl*II and cloned into *Kpn*I/*Bgl*II -digested pBBR1MCS-5 to create pBBR1MCS-6. The full *ttg2* operon (3650 bp) from *P. aeruginosa* PAO1 was cloned separately into pBBR1MCS-5 or pBBR1MCS-6 as described in Table S4.

**RNA extraction and real-time quantitative PCR.** Early-log-phase cultures of the mutant and complemented strains grown on LB were adjusted to obtain a suspension of approximately $10^9$ cell $ml^{-1}$. 0.5 ml of the bacterial suspension were treated with 1 ml of the RNAprotect Reagent (Qiagen) according to the manufacturer's instructions for stabilization of the RNA molecules. Total RNA was extracted by using RNeasy Mini Kit (Qiagen) with on-column DNase digestion (RNase-Free DNase set, Qiagen) to remove contaminating DNA. Removal of DNA was confirmed by performing PCR using an aliquot of the DNase-treated RNA as a template. Reverse transcription reactions were carried out in 20-µl volume containing 0.1 µg RNA, random primers, and the buffer and enzyme components of the Maxima First Strand cDNA Synthesis Kit for RT-qPCR kit (Thermo Fisher Scientific) according to the supplied protocol. Initial PCR amplifications for the expression of *ttg*2 operon-specific mRNAs were performed on the cDNA templates from the parental strains, the Δ*ttg*2 mutants, and the complements to confirm the loss of gene expression in the Δ*ttg*2 deletion mutants and recovery in the complemented strains. Real-time qPCR analysis was carried out on the CFX96 machine at the genomics core facility of Universitat Autònoma de Barcelona using a PCR master mix containing SYBR green dye. The sequences of the *ttg*2D primers used in the real-time qPCR (Int-*ttg*2D-U and Int-*ttg*2D-L) are given in Table S4. These primers amplify a 215-bp conserved portion of the *ttg*2D gene. Relative gene expression comparisons were obtained through the $\Delta\Delta C_T$ method (CFX Manager software) by normalizing the mean cycle threshold of the investigated transcript to the housekeeping gene *rpsL* with primers rpsL-F; GCAAGCGCATGGTCGACAAGA) and rpsL-R; CGCTGTGCTCTTGCAGGTTGTGA (amplicon size 201bp).

**Outer membrane permeabilization assay.** The NPN (1-*N*-phenylnaphthylamine) uptake assay was done according to Loh *et al.* [33], with modifications. Briefly, overnight cultures of the different strains in MHB were subcultured into the same medium and grown to mid-logarithmic phase. Appropriate antibiotic was added to growth plasmid-bearing strains. Cells were washed with 10 mM sodium HEPES (pH 7.2), and then resuspended at a final $OD_{550nm}$ of 1.0 in the same buffer supplemented with a 5 µM CCCP (carbonyl cyanide *m*-chlorophenylhydrazone). CCCP was added to block energized secretion of NPN and to prevent a decline in fluorescence during the assay [33]. 50 µL of cell suspension was pipetted into a quartz cuvette containing NPN (10 µM final concentration), and as test substances either EDTA (0.2 mM final concentration) or colistin (10 µg $ml^{-1}$ final concentration) to a total volume of 100 µl. Fluorescence was monitored by a Cari Eclipse spectrophotometer (Variant, Inc., Palo Alto, C.A) at excitation and emission wavelengths of 340 nm and 415 nm, respectively. Control experiments without added cells or without colistin or EDTA were also performed. Each assay was performed at least three times.

**Analysis of antimicrobial susceptibilities.** Minimal inhibitory concentration (MIC) to antibiotics of several classes including front-line antipseudomonal drugs, namely imipenem, meropenem, amikacin, tobramycin, ceftazidime, piperacillin/tazobactam, ciprofloxacin, levofloxacin and colistin, was determined by the broth microdilution method

(BMD) or by Etest. The BMD method was performed on cation adjusted MH broth (CAMHB) as recommended [34, 35]. Bacteria were first grown overnight in CAMHB using CLSI-recommended incubation conditions and the antibiotics were serially diluted twofold across the 96 well plates. After that, 100 µl of a bacterial suspension diluted to $5x10^5$ CFU/ml in CAMHB was added to the wells containing the antibiotic dilutions. The 96-well plates were incubated for 20 h at 37°C before developing by visual inspection or with the resazurin dye [36]. MIC was defined as the lowest concentration of the antibiotic (in µg $ml^{-1}$) that prevented visual growth. As the LESB58 isolate is considered a slower growing strain compared to laboratory strain PAO1 [37], incubation for bacterial growth was extended to 48 h instead of 24 h for some protocols. MICs were confirmed by two or three independent replicates, and MIC values 2-fold or greater than that of the control were considered significant. For Etest a 0.5 McFarland suspension was used to create a confluent lawn of microbial growth in 150-mm Mueller-Hinton (MH) agar plates. The MIC values were determined according to the Etest reading guide after 18 h incubation at 37°C. MICs were determined as the concentration at which the zone of inhibition intersected the Etest strip.

**Tolerance to organic solvents, and SDS/EDTA**

The approach to assess solvent tolerance involved overlaying solvent (100%) onto LBMg (LB medium supplemented with 10 mM $MgCl_2$) agar plates (55 mm glass plates) inoculated with bacteria as previously described [38]. Briefly, late exponential phase LB broth cultures were diluted into the same medium to yield a suspension of approximately $10^7$ cells $ml^{-1}$. A 5-µl aliquot of the cell suspension was spread over the surface of an LBMg agar plate in duplicate and allowed to dry before being overlaid with an organic solvent to a thickness of 3 mm. The plates were sealed and growth was assessed following incubation at 37°C for 24-48 h. Wild-type PAO1 was unable to grow in modified LB agar in the presence of toluene or *n*-hexane (data not shown), but it grew well in LBMg plates overlaid with *p*-xylene. MIC assays were conducted in 96 well plates to determine the bacterial sensitivity to SDS and/or EDTA. The MIC values were defined as the lowest substance concentration that inhibited 80% of growth (based on OD measurements) in comparison to the growth control.

**Biofilm formation.** Biofilm formation was assessed as previously described [39] with the following modifications. Sterile 96-well flat bottom polystyrene non-treated plates (BrandTech 781662) were used. Two-hundred microliters of overnight cultures adjusted to an $OD_{550nm}$ of 0.1 were incubated in LB broth or LB supplemented with 0.05 mM EDTA for

24 hours at 37°C. Cells were washed three times with water, fixed at 60°C for 1 h and stained during 15 minutes with 200 µl of 0.1% crystal violet. The dye was discarded and the plate was rinsed in standing water and allowed to dry for 30 min at 37°C. Crystal violet was dissolved in 250 µL 95% ethanol for 15 min, and the OD of the extracted dye was measured at 550 nm. Biofilm formation was normalized by cell growth and reported as relative biofilm formation. A one-way ANOVA with Tukey's multiple comparison test (GraphPad Prism 6.0) was used to determine the significance of the data between groups.

**Protein *in-silico* analysis and bioinformatics tools.** Protein sequences were analyzed using BLAST, PSI-BLAST and CDD within NCBI (http://ncbi.nlm.nih.gov/) and PsortB (http://psort.nibb.ac.jp). Known 3D structures for *P. aeruginosa* Ttg2D orthologs were downloaded from PDB (https://www.rcsb.org/). Structures with accession numbers 2QGU (*R. solanacearum*) and 4FCZ (*Pseudomonas putida*) were deposited at the PDB by the Northeast Structural Genomics Consortium (NESG) without any associated publications. Structures 5UWA (*E. coli*) and 5UWB (re-refined coordinates for 4FCZ) were submitted to the PDB by Ekiert *et al.* [2].

Homologous sequences were obtained from a set of representative proteomes at Pfam database (https://pfam.xfam.org/). First, the full sequences of the PF05494 (MlaC family) RP15 group members were downloaded, and Ttg2D orthologs in Gram-negative organisms were then selected from reciprocal best hits. In addition, sequences from unclassified bacteria were removed from the list, totaling 151 representative protein sequences for further analysis. Multiple sequence alignment using a hidden Markov model (HMM) profile was generated with the hmmalign program of the hmmer3 package (http://hmmer.org/), and sequence logos were generated with Weblogo3 [40]. For phylogenetic analysis, an unrooted maximum likelihood tree was reconstructed using the best model of evolution on MEGA 7 [41], based on amino acid sequences. Phylogenetic tree was visualized and annotated using the interactive web platform iTOL v3 [42].

Superposition of the *P. aeruginosa* Ttg2D and 2QGU structures was performed by aligning both sequences to the Pfam HMM profile for the MlaC family with the hmmaling program from the hmmer3 package, and using the resulting alignment to superpose both structures with Profit (http://www.bioinf.org.uk/programs/profit). In order to avoid superposing residues from the C-terminal bended helix, only residues 1 to 175 from Ttg2D and 29 to 199 from 2QGU where used. Superposition of the Ttg2D structure to representative structures of substrate-binding protein subclusters [1] was carried out with the Mammoth program [43]. Structural domains where assigned by transferring those defined for PDB structure 2QGU in CATH database [44]. The search for similar structures in the whole PDB database was performed using Dali server [45].
